# Inferring assembly-curving trends of bacterial micro-compartment shell hexamers from crystal structure arrangements

**DOI:** 10.1101/2022.10.21.512534

**Authors:** Luis F. Garcia-Alles, Vanessa Soldan, Irma Custovic, Sara Castaño-Cerezo, Stéphanie Balor, Miguel Fuentes-Cabrera, Gilles Truan, Eric Lesniewska, David Reguera

**Affiliations:** TBI, Université de Toulouse, CNRS, INRAE, INSA, 135 Avenue de Rangueil, 31077 Toulouse, France; METi, Centre de Biologie Intégrative, Université de Toulouse, CNRS, UPS, 31062, Toulouse, France; ICB UMR CNRS 6303, University of Bourgogne Franche-Comte, Dijon, F-21000, France; Center for Nanophase Materials Sciences, Oak Ridge National Laboratory, Oak Ridge, Tennessee 37831, United States; Departament de Física de la Matèria Condensada, Universitat de Barcelona, Martí i Franquès 1, 08028, Barcelona, Spain; Universitat de Barcelona, Institute of Complex Systems (UBICS), Martí i Franquès 1, 08028, Barcelona, Spain

**Keywords:** bacterial micro-compartments, shell, hexamer, assembly curvature, crystal structure, microscopy, nanotube

## Abstract

Bacterial microcompartments (BMC) are complex macromolecular assemblies that participate to varied chemical processes in about one fourth of bacterial species. BMC-encapsulated enzymatic activities are segregated from other cell contents by means of semipermeable shells, justifying why BMC are viewed as prototype nano-reactors for biotechnological applications. Herein, we undertook a comparative study of trends of self-assembly of BMC hexamers (BMC-H), the most abundant shell constituents. Published and new microscopy data show that some BMC-H, like *β*-carboxysomal CcmK, tend to assemble flat whereas other BMC-H often build curved-implying objects. Inspection of available crystal structures presenting BMC-H in tiled arrangements permitted to identify two major assembly modes with a striking connection with experimental trends. All-atom molecular dynamics (MD) supported that BMC-H bending is triggered robustly only from the disposition adopted by BMC-H that form curved objects experimentally, conducting to almost identical arrangements to those found in structures of recomposed BMC shells. Simulations on ensembles of planar-behaving hexamers, which were previously reconfigured to comply with such disposition, confirmed that bending is defined by assembly details, rather than by BMC-H identity. Finally, although no common atomic determinants could be identified as responsible of BMC-H spontaneous curvature, an inter-hexamer ionic pair was pinpointed as contributor to hold a subset of BMC-H in low bending dispositions. These results are expected to improve our understanding of the variable mechanisms of biogenesis characterized for BMC, and of possible strategies to regulate BMC size and shape.

## INTRODUCTION

Bacterial micro-compartments (BMC) are complex macromolecular protein assemblies that confine specialized enzymatic activities within shells and participate to processes like carbon fixation in cyanobacteria or metabolite degradation related to bacterial growth and pathogenesis ^1^. According to a recent genomic survey, about 70 BMC types can be found in nature, all differing by sort of cargo contents ^2^. In contrast, all shell protomers belong to two structural families. The first one adopts the Pfam 00936 fold, which associate as hexamers (BMC-H) or trimers of bidomain proteins (BMC-T). These proteins can be further sub-classified depending on structural details such as for instance the permutation of secondary elements or the capacity to further dimerize ^3, 4^. Members of the second family (Pfam 03319) assemble as pentamers (BMC-P) and occupy shell vertices ^5^. Accordingly, BMC-P are stoichiometrically very minor in shells.

The structure of reconstituted minimalist shells proved that BMC-H are endowed with a considerable assembly versatility, being capable of establishing contacts with themselves in at least two different dispositions (planar or bent), but also with BMC-T and BMC-P shell partners ^5^. Noteworthy, the same set of residues basically ensured interactions with all different partners, irrespective of the precise local symmetry environment, something that is reminiscent of viral capsids, where a single protein often occupies different structural environments. Yet, BMC shells were demonstrated in atomic-force microscopy (AFM) nano-indentation experiments to be considerably less rigid than capsids or encapsulins ^6^.

Despite impressive advances in the structural characterization of BMC, our understanding of shell assembly is progressing slowly. A major milestone was the characterization of the biogenesis of *β*-carboxysomes from *Synechococcus elongatus* PCC 7942 (*Syn7942*) ^7^. Colocalization experiments with fluorescently-labelled RuBisCO and shell proteins evidenced that *β*-carboxysome assembly is a two-step process, shells appearing only after the emergence of pre-organized procarboxysome “grains” of coalesced RuBisCO, carbonic anhydrase and scaffolding proteins (*cargo-first* mechanism, see below). The presence of less organized/compact encapsulated contents in other BMC types, when compared to hexadecameric RuBisCO ^8^, and the observation of compartment formation in the absence of cognate cargo ^9, 10^, pointed to the existence of other assembly pathways for other BMC types. In line with this, a recent study on Pdu BMC biogenesis proved that *cargo-first* and *shell-first* assembly pathways are both feasible, even when cargo-enzymes coalesce following a hierarchical organization mode ^11^.

A compelling understanding of assembly is being provided by theoretical simulations using coarse-grained (CG) models to describe simplified cocktails of shell components, cargo proteins and even scaffolding factors ^12, 13^. These studies indicated that assembly pathway, as well as the morphology and cargo-loading extent will be function of the precise balance of interaction strengths between the components and of their stoichiometries. For instance, strong scaffold-mediated cargo-cargo interactions would favour two-step mechanisms, whereas weaker interactions would lead to concomitant scaffold-cargo coalescence and shell assembly. These studies also highlighted that, albeit BMC size is relatively insensitive to hexamer-hexamer or scaffold-hexamer affinities, it was critically impacted by the assignment of spontaneous shell curvature and scaffold length ^13^. Curvature would result from an imbalance of attractive and repulsive forces established above and below the planes of each interacting pair of hexamers.

Although expected to be critical in driving BMC closure, studies of spontaneous curvature are scarce. To the best of our knowledge, only the assembly of PduA was investigated with this intention ^14^. All-atom molecular dynamics (MD) performed on a pair of interacting hexamers revealed a preference to remain planar. Notwithstanding, coarse-grained simulations were parametrized with 25° curvature between hexamers, a choice based on angles observed in the cryo-electron microscopy (EM) structure of *Hal. ochraceum* (HO) BMC shells ^5^. Globally, it is still unknown whether BMC-H are endowed with properties required to induce spontaneous shell curvature or not.

In this study we present experimental evidences that support the existence of two major BMC-H assembly behaviors, depending on the preference to form curved-implying or flat objects, and the possibility that this could be inferred from the kind of organization adopted in crystal structures exhibiting piled planar subunit arrangements. Indeed, all-atom MD performed on hexamer ensembles extracted from these structures closely reproduced experimental trends. Thus, BMC-H that form curved objects and display the first organization mode in crystals, bent rapidly and reliably, evolving towards similar dispositions to those characterized in objects formed by purified proteins belonging to this group ^15^ or in structures of minimalist reconstituted BMC shells ^5, 16^. On the contrary, hexamers adopting other assembly types were more reluctant to bend. Although interaction energy profiles along the hexamer interface were shallow, indicating that assembly is cooperative, an ionic interaction was found to contribute to hold a BMC-H subset away from the disposition that precedes shell formation.

## RESULTS

### BMC-H propensity to assemble

Individual BMC-H have been shown to form varied structures *in vivo*, generally after recombinant expression in *E. coli* (a compilation of literature data is given in Table S1). EM and AFM studies on purified proteins from this family also highlighted an extraordinary assembly plasticity. Thus, apart from predicted 2D organizations, filaments/nanowires, nanotubes, swill-rolls and even spherical structures could be characterized. With the intention to contribute more data and complete the vision, we programmed the characterization of several purified oligohistidine-tagged BMC-H, which were not covered in previous studies, or only partly: CcmK1, CcmK2 and CcmK4 from *Syn6803*, CcmK2 and CcmK4 from *Syn7942*, CsoS1A from *H. neapolitanus*, PduA and PduJ from *S. enterica*, EutM from *E. coli*, BMC-H from *H. ochraceum* and RMM-H from *M. smegmatis* [referred hereafter with the notation: Protein name^*species abbreviation*^ (e.g. CcmK1^*6803*^)].

Two types of data permitted to get a glance of BMC-H relative assembly potential. First, the alteration of cell size induced upon BMC-H overexpression was assayed using phase contrast microscopy (Fig. S1A). When compared to controls, a significant number of PduA^*Sent*^ and PduJ^*Sent*^ expressing cells exhibited increased length, something that also occurred at lower extent with RMM-H^*Msm*^ and BMC-H^*Hoch*^. Size alterations were considerably weaker than those reported ^17^, something possibly related to the different induction methods. Expression of *Syn6803* proteins also led to a moderate increase of length, in comparison to empty pET15b controls. Although data collected for CcmK3^*6803*^, an insoluble protein when expressed in *E. coli*, invite to caution, collectively these results suggest that nanotubes or 2D-assemblies form inside cells, as previously reported for PduA and PduJ ^17^ or other assembly-prone proteins ^18^. In the second approach, BMC-H remaining soluble after IPTG induction was compared with amounts of potential assemblies retained in pellets. The latter was inferred using an experimental setup that pursued the solubilization by disassembly with a chaotropic agent, urea, at sufficiently low concentrations as to avoid the recovery of misfolded proteins forming inclusion bodies (IB). Experiments with CcmK3^*6803*^, an insoluble protein that only becomes soluble when co-expressed with CcmK4 ^19^, indicated that resolubilization of IB required at least 2 M urea concentrations.

With the exception of C-ter Histagged CsoS1A^*Hneap*^, all other tested BMC-H were correctly expressed in *E. coli* (Fig S1B). Globally, BMC-H distributed between soluble fractions (SF) and the urea solubilized fraction (USF), the latter being most prominent for most of proteins. PduA^*Sent*^, CcmK1^*6803*^, CcmK4^*6803*^ and CcmK4^*7942*^ were almost exclusively recovered in urea fractions. On the opposite side, EutM^*Ecol*^ and CcmK2^*6803*^ were more abundant in soluble fractions. Noteworthy, urea purified fractions (UPF) remained soluble after removal of urea and gave similar results in AFM and transmission EM (TEM) as the PF. The pentameric CcmL^*6803*^ was also found at relatively high amounts in urea fractions, suggesting potential assembly, as reported ^20^.

Overall, these data indicated a common strong propensity of BMC-H to form big macro-molecular entities during expression inside bacteria.

### Two general BMC-H assembly behaviors

To complete published data compiled in Table S1, we investigated the assembly of CcmK from *Syn6803* and *Syn7942* inside *E. coli* by TEM (Fig S2). Only CcmK1^*6803*^ and CcmK3^*6803*^ images differed from those obtained for the control strain (empty vector). Although unclear, stripped motifs might occur with CcmK1^*6803*^. With CcmK3^*6803*^, homogenous spherical patches suggestive of IB were often present at cellular poles. The lack of noticeable observations for CcmK proteins contrasts with nanotubes, filaments and spherical structures reported for many other BMC-H (Table S1).

Assembly potential was also assessed by AFM and TEM taking purified proteins conditioned at comparable concentrations in the presence of a phosphate buffer. 2D flat-organizations were recurrently imaged by AFM with all CcmK proteins (see Fig. S3A-F), although CcmK1^*6803*^ gave rise to the reported honeycomb mosaics consisting of laterally-collapsed cup-like motifs^21^. Less clear-cut and difficult to reproduce data were obtained for the rest of BMC-H, pointing to difficulties expected if rounded objects formed, since these would establish weak contacts with the planar mica support. 2D patches were seen rarely with PduJ^*Sent*^ (Fig. S3M,N), and with BMC-H^*Hoch*^ (Fig. S3O), in agreement with reported data collected for the untagged protein ^22^. With PduA^*Sent*^ or EutM^*Ecol*^, mica was sometimes covered with material that attained relatively homogenous heights (less than 6 nm, Fig S3J, S3P). Notwithstanding, patches remained small, suggesting difficulties to grow in 2D. Spheroids insinuated in occasions with CsoS1A^*Hneap*^ and PduA^*Sent*^ (Fig. S3G,I,K). Similar spherical entities were also imaged as protrusions/incrustations with RMM-H^*Msm*^ or EutM^*Ecol*^ (Fig. S3Q). Linear arrangements evocative of bundles of nanotubes were monitored for PduA^*Sent*^ in a buffer supplemented with MgCl_2_ (Fig. S3L), but could not be reproduced in independent experiments. Among all CcmK imaged by TEM, assemblies were only evident for CcmK1^*6803*^ (Fig. S4A-B): flat 2D patches with irregular edges, occasionally also polygonal-like tiles. Roughness was noticed inside some assemblies, which could reflect abovementioned honeycomb mosaics. Flat assemblies also occurred with PduA^*Sent*^ (Fig. S4F-G), PduJ^*Sent*^ (Fig. S4H-I) or BMC-H^*Hoch*^ (Fig. S4L-M). Hexamers were more evident with the latter, in virtue of spacing close to the 7.5 nm distances reported before ^23^. Patches had remarkable sharp edges and almost polygonal shapes in the case of the two Pdu proteins. However, the most recurrent structures imaged for PduA and PduJ were nanotubes, in agreement with the literature (see Table S1). Frequently appearing straight, with 22-30 nm diameters in the case of PduA, wider with PduJ, their darker interior suggested the access of TEM contrast agent to their lumen. PduA and, especially, PduJ nanotubes stuck sometimes on 2D tile-edges (e.g. Fig. S4I), suggesting a potential 2D-growth mechanism. Indeed, although not studied here, we noticed that the probability of observing flat tiles with PduA augmented with prolonged incubations in saline phosphate buffer. For PduJ, some images suggested the possibility that large nanotubes, apparently opened along the longitudinal axis, were in fact partly rolled 2D-assemblies (e.g. structure pointed by the arrow in Fig. S4H). Spheroids and narrow curved filaments/nanowires (16-20 nm diameters) were sporadically detected with PduA^*Sent*^, CsoS1A^*Hneap*^ (detected in only one out of three independent preparations, Fig. S4C-E) and EutM^*Ecol*^ (Fig. S4J-K). Indeed, EutM^*Ecol*^ more often resulted in skinny and aggregated material without evident organization, as judged by FT treatments. Amorphous fibers were seen in few occasions for RMM-H^*Msm*^, which might be reminiscent of bundles of nanotubes observed at significantly higher RMM concentrations ^24^. Formation of varied types of assemblies was confirmed in cryo-EM experiments, both for PduA and PduJ (Fig. S5A-F), whereas large and dense nanotubes with rough edges were observed with CsoS1A (Fig. S5G-I).

These data and reported observations, apart from highlighting an extraordinary assembly plasticity, point to the manifestation of two major trends: some proteins are prone to build curved-implying objects (PduA^*Sent*^, CsoS1A^*Hneap*^, RMM-H^*Msm*^ and possibly PduJ^*Sent*^); *β*-carboxysome CcmK proteins and possibly BMC-H^*Hoch*^ more easily organize as (quasi)flat assemblies. Less clear-cut, EutM^*Ecol*^ might figure among curving BMC-H, according to published data.

### Different 2D-assembly modes identified in BMC-H crystals

Our intention was to investigate BMC-H assembly behavior by MD simulations taking advantage of crystallographic data. By the time of the realization of this study, there existed about 60 BMC-H structures deposited in the Protein Databank (plus 8 entries from reconstructed shells published in the course of this work, see Table S2). Thirty-four structures were from wild-type (WT) proteins. Within this group, we focused our work on the 18 crystal structures that displayed hexamers organized as piled 2D layers (see below).

These structures could be categorized in four groups according to the BMC-H disposition, which differed by lateral displacements and distances between interacting hexamers (Fig. 1 and Table S2). The first assembly type (hereafter called *Ass-A*) is characterized by a short distance between the two key Lys residues from interacting hexamers. A representative case is the PduA^*Sent*^ 3NGK structure, with measured 7.2-8.4 Å distances between Lys26 Cα from hexamer counterparts (Fig. 1A). *Ass-A* is the only assembly mode observed for WT PduA^*Sent*^, CsoS1A^*Hneap*^, CsoS1C^*Hneap*^ and BMC-H^*Ahyd*^ (Fig. 1B). *Ass-A* also reflects closely the disposition of BMC-H noticed in all reconstituted shells. The second assembly mode (*Ass-B*) is adopted by all CcmK proteins, EutM^*Ecol*^ and BMC-H^*Hoch*^ (Fig. 1C-D). Here, the inter-lysine distance is considerably longer (14.9-17.6 Å). An intermediate organization (*Ass-C*), with distances of about 10 Å, is seen in CcmK2^*6803*^ 3DNC and CcmK4^*7942*^4OX6 structures, whereas CcmK2^*7942*^ 4OX7 is the only case displaying a fourth assembly type (*Ass-D*). Assembly occurrence did not seem to be defined by crystallization conditions. Thus, the organization mode is reproduced in crystals of the same protein prepared under variable conditions (e.g. *Ass-A* for structures 2EWH and 2G13 from CsoS1A^*Sent*^, or in the 3H8Y structure of the close CsoS1C^*Sent*^ homolog; *Ass-B* for 3BN4 or 3DN9 structures of CcmK1^*6803*^ or 3MPW and 3MPY from EutM^*Ecol*^).

**Figure 1.**
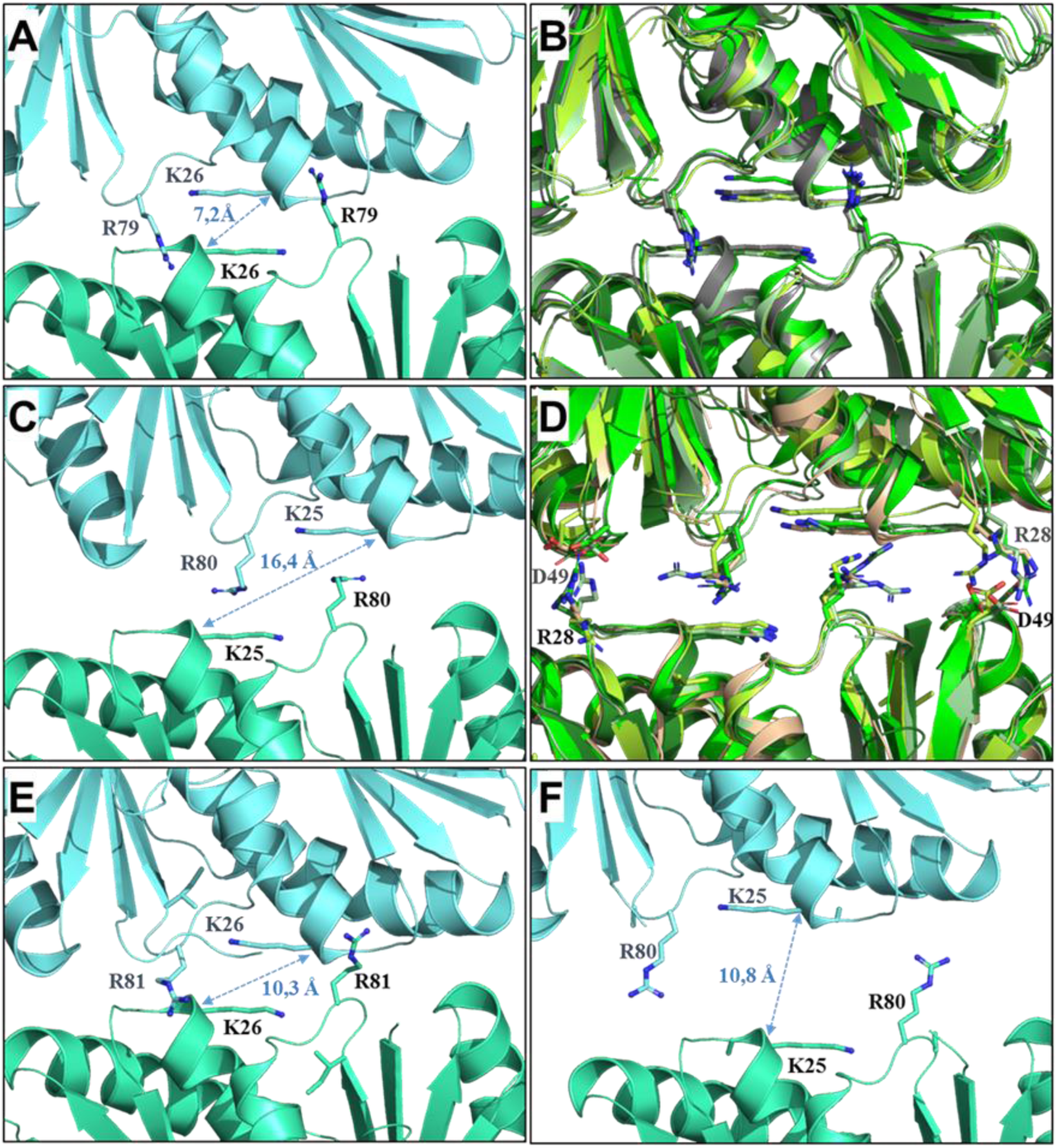
Four general 2D-arrangements in crystals of BMC-H. One representative case for each of the four organizations is presented in next panels: *A*, PduA^*Sent*^ (3NGK) for the *Ass-A* organizaion; *C*, CcmKl^*6803*^ (3BN4) for *Ass-B; E*, CcmK4^*7942*^ (40×6) for *Ass-C* and *F*, CcmK2 ^*7942*^ (40×7) the unique member displaying the *Ass-D* arrangement. A close-up view of the inter-hexamer interface is shown with each ribbon-represented hexamers colored cyan or blue marine. Side-chains of key Lys and Arg are shown as sticks, with nitrogen atoms in deep blue. Distances are measured between alpha carbons of the two Lys. Panel B presents a view of structures with proteins adopting *Ass-A* dispositions: PduA (3NGK) in grey, PduJ (5D6V) with restored K25 in green, CsoSlA (2G13) in pale green, CsoSlA (3H8Y) in limon and BMC-H^*Ahyd*^ (4QIV) in forest green. The view was generated after super-imposition on main-chain atoms from the hexamer seen at the bottom. A similar image was prepared in panel D for proteins of Ass-8 group: CcmKl^*6803*^ (3BN4) in green, CcmK2^*6803*^ (2A1B) in pale green, CcmK4^*6803*^ (6SCR) in limon, EutM^*Ecol*^ (3MPW) in forest green and BMC-H^*Hoch*^ (5DJB) in wheat. Ionic interactions between Arg28 and Asp49 of CcmKl^*6803*^, or corresponding residues, are established in Ass-B organizations.

Globally, a parallel seem to exist between experimental assembly behavior and the way BMC-H tile in crystals. *Ass-A* would be the preferred crystallization mode for proteins endowed with curving propensity, whereas flat-behaving BMC-H would mostly adopt *Ass-B* arrangements.

### Variable curving trends supported by all-atom molecular dynamics

The last hypothesis was challenged by means of all-atom MD. We followed the approach described in a previous study ^21^, in which MD runs were launched on ensembles of three interacting hexamers, extracted from crystal structures showing tiled BMC-H (PDB codes indicated in Table S3). Tri-hexameric ensembles were selected as best compromise to describe the situation in BMC shells while keeping reasonable computational costs. After energy minimization, MD were run (AMBER forcefield) with proteins embedded in explicit hydration boxes under quasi-physiological conditions (see M&M). All different assembly types mentioned in the previous section were covered. Hexamer tilting and bending angles were monitored for intermediate structures extracted in the course of each trajectory (250 ps snapshots), as well as inter-hexamer distances calculated from the hexamers center of mass in the MD average structure. Rather than long single simulations, we opted by performing several independent simulations on each case (20 ns each), which differed by the attribution of random initial atom velocities. This parameter is known to impact MD trajectories more profoundly than time length ^25^, something that we indeed confirmed (see below).

MD behavior seemed to be defined by the kind of assembly. Thus, strong and reproducible bending was noticed for all *Ass-A*-deriving cases, tilting remaining close to values measured for the crystal (Fig. 2, Fig. S6 - S7, and Table S3). For instance, PduA^*Sent*^ and CsoS1A^*Hneap*^ (2G13) bent by about −25°, something reproduced in 4 independent runs. Among *Ass-A* cases, the less pronounced effect occurred for PduJ^*Sent*^ (starting ensemble taken from the K25A mutant 5D6V entry, after manual reintroduction of interfacial K25 side-chains). As expected for bending, inter-hexamer separation decreased slightly (not to be confused with edge to edge inter-hexamer separation) (Table S3). Data dispersion for all *Ass-A* BMC-H was low, even smaller than those measured for EutL^*Ecol*^, a BMC-T that basically remained flat in two MD runs. Induction of curvature was rapid, reaching dispositions close to the MD average during the first nanosecond (Fig. 2D), explaining the narrow data dispersion. Importantly, negative bending values corresponded to the orientation described for full BMC shells. Indeed, root-mean-square deviations (RMSD) of 1.3 Å were measured (on 1392 backbone atoms) when comparing interacting pairs of monomers in the average structure of two independent MD PduA^*Sent*^ (3NGK) with corresponding interacting monomers of bent BMC-H in the *H. ochraceum* BMC shell structure (5V74). Worth-mentioning, the results were reproduced using CHARMM as forcefield in similar independent simulations launched on PduA^*Sent*^ (average bending of −23°) and CsoS1A^*Hneap*^ (−16°).

**Figure 2.**
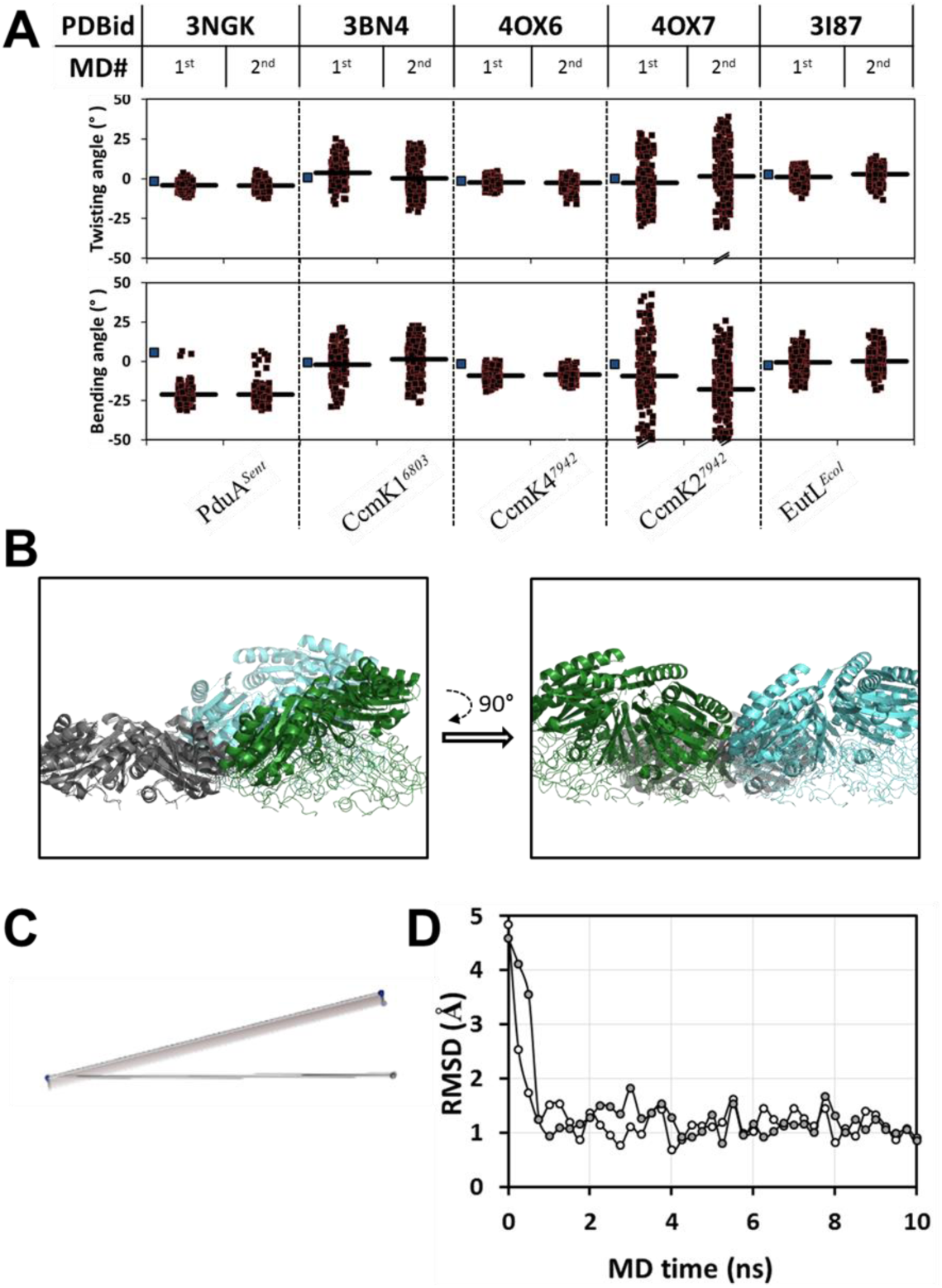
MD simulations on BMC-H tri-hexamers. *A*, Tilting and bending angles calculated from all-atom MD trajectories of ensembles of three BMC-H, originally positioned as in published crystal structures (indicated PDBid codes). Each point (black square) corresponds to a snapshot. Two independent 20 ns simulations are presented separatedly (1^st^ and 2^nd^). Angles were calculated from mainchain alpha carbon (Ca) atoms of selected residues in interfacial monomers from each hexamer (see M&M). One representative case from each kind of assembly type is shown (PduA^*sent*^ for *Ass-A*, CcmKl^*6803*^ for *Ass-B*, CcmK4^*6803*^ for *Ass-C* and CcmK2^*7942*^ for *Ass-D*). With the latter, values for some snapshot were out of the graph. For comparison, data for EutL^Ecol^ BMC-T are presented in the last two columns. Thick traces are the mean value for the MD run. Blue squares preceding each case represent angles measured in the original crystal structure.B, Comparison of the average structure of the first MD simulation with PduA^Sent^ (cartoon) with the disposition at time 0 (thin traces). The two structures were superposed on backbone atoms of one hexamer. Two orthogonal views are shown. *C*, The same comparison is illustrated with planes that were elaborated from the coordinates of the center of mass (COM) of each hexamer. The plane composed by spheres in grey represents the starting structure. Spheres in blue are for the MDs average structure. The view is approximately seen as in panel B. Consequently, the traverse view of the plane in the crystal structure results in a flat trace. Bending during the MDs induces the trace to displace up or down. *D*, Assembly evolution in the course of PduA^sent^ MD simulations. Plotted are the RMSD values calculated when the coordinates of backbone atoms from each snapshot structure (0,25ns steps) are compared to the MD average structure (empty circles for first MD, gray for the second run).

Assembly fate was more uncertain for non-*Ass-A* organizations. Bending and tilting were variable (Fig. 3), the reproducibility between runs being considerably poorer than with *Ass-A*, as illustrate strong data dispersion (Table S3). With some cases, the tri-hexamer remained close to flat independently of the run [e.g. CcmK1^*6803*^ (3BN4) or CcmK4^*7942*^]. But for most other cases, behavior was instable, even conducting to inversions of bending orientation (e.g. EutM^*Ecol*^ or BMC-H^*Hoch*^). As possible exception, CcmK2^*6803*^ (3DNC) evolved from its *Ass-C* dispositions in the two runs much like *Ass-A* assemblies. Yet, the overall trend for non-*Ass-A* dispositions seemed to be to remain planar.

**Figure 3.**
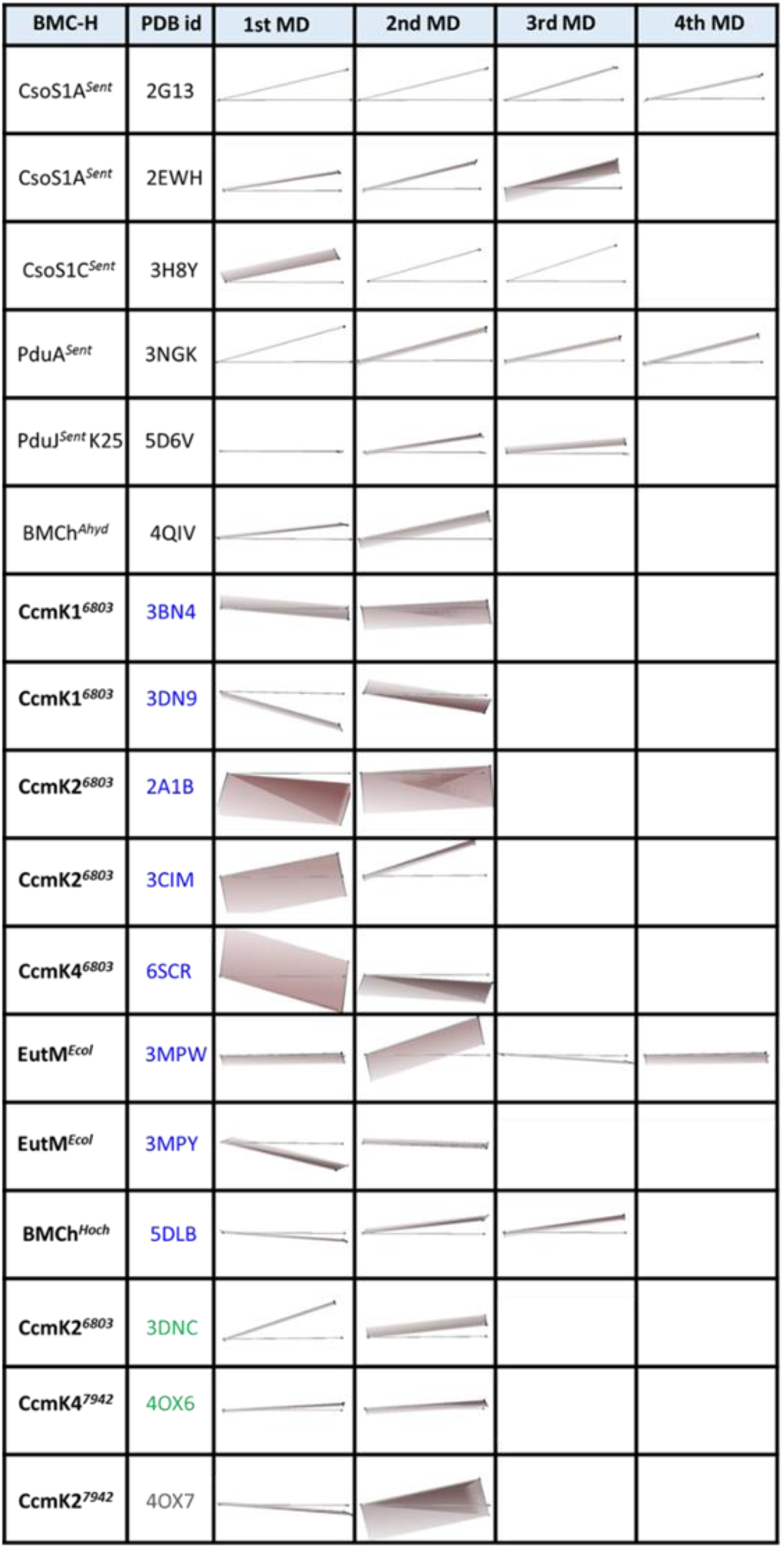
Comparison of average structure from MD simulations on BMC tri-hexamer ensembles with starting crystal structure. Average structure generated for all snapshot structures extracted from each MD simulation compared with the disposition at time 0. The two structures were superimposed on backbone atom coordinates of one of the hexamers (sphere representing the hexamer centers of mass (COM) most to the left). Each assembly is represented by the plane that passes by the COM spheres of the three hexamers, which are in grey for the starting structure, in blue for the MDs average structure (see Fig. 2C). PDB codes are colored to indicate the type of organization: black for *Ass-A* arrangements, blue for *Ass-B*, green for *Ass-C* and grey for *Ass-D*.

### Theoretical behavior of shell assembled BMC-H substructures

The possibility that *Ass-A* configurations were responsible of BMC-H curving was evaluated in simulations launched on ensembles of 2 interacting BMC-H extracted from BMC shells (tri-hexamers are not found so far in shells), since BMC-H always adopted *Ass-A*-like dispositions in characterized structures (Table S2). The two hexamer configurations observed for the BMC-H^*Hoch*^ 5V74 shell were assayed^5^, which correspond to either bent or flat organizations, as well as an ensemble of two bent CcmK^*7418*^ hexamers from the 6OWF shell structure^16^.

MDs trajectories clearly showed a preference for curved dispositions: bent BMC-H^*Hoch*^ or CcmK^*7418*^ remained close to the starting angle, while the planar BMC-H^*Hoch*^ evolved towards a curved configuration (Fig. S7).

### Atomic determinants for triggering BMC-H bending

Our most challenging aim was the identification of atomic determinants implied in curvature. We selected for such study PduA^*Sent*^, because of its demonstrated experimental trend to form nanotubes and robust MD behavior. First, we sought to establish key interactors that clamp hexamers together. Two analytical approaches were followed: i) side-chain RMSD with regard to the MD average structure were monitored over the MD run (Fig. S8 and S9). For each residue, the different 18 positions in the tri-hexamer were plotted together. A clamping residue was expected to result in a relatively fixed conformation, and thus in lower RMSD, but only when located at the interface. In that manner, Lys26 and Arg79 always occurred with lowest RMSD at interfaces, for two independent MDs. Interfacial Glu19, Asp22, Asn29, Pro78 and His81 were often, but not always, lowest; ii) the contribution to the interaction energy of each residue was evaluated. The snapshot with lowest RMSD to the average structure of each MD was selected and energy-minimized. We considered as contributor to the interaction those residues that resulted in maximal interval of values, when comparing values for the 18 monomers (Fig. S10A), and at the same time presented highest stabilization when at the interface, when compared to exposed positions (Fig. S10B). In that manner, Lys26, Arg79 were again confirmed to be key contributors. Surprisingly, the only additional identified residue was Ser27.

The role of above-pinpointed residues for assembly fate was investigated by *in silico* MD of single-residue alanine mutants. We extended the exploration to other interfacial residues to anticipate unforeseen possibilities, also to positions known to establish contacts between hexamers in characterized BMC shells, such as Asn67 that replaces Arg66 of CcmK^*7418* 16^. In total, the next 21 residues were scanned: K12, E19, D22, K26, S27, N29, R48, D50, V51, K55, D59, R66, N67, H75, P78, R79, H81, T82, D83, E85 and K86 (mutations were introduced 6 or 9 times in the tri-hexamer, depending on whether the residue was close to the center or edge of the interface, respectively). Notably, curvature was only impacted by the K26A mutation (Fig. S11), which completely abolished bending and led to 1-2 Å larger hexamer separation. The result was reproduced in four independent 20 ns MDs (only two shown). All other mutations were without effect, including the R79A. None of 6 side-chains tested replacing the interfacial K26 was able to restore bending (Fig. S12), a result that agrees with the full conservation of this residue among BMC-H ^3^.

The importance of the two key interfacial residues was evaluated in other BMC-H (Fig. S13). CsoS1A^*Hneap*^ behavior was not perturbed by either K29A or R83A mutations. Bending extent of K25A BMC-H^*Ahyd*^ diminished slightly, depending on MD run, whereas the R78A mutation was again without consequence.

### Multiple energy minima in lateral contacts between planar BMC-H

MD results combined with the structures of all recomposed shells (Table S2) concur to prove that *Ass-A* is the ready-to-curve disposition. That most other BMC-H tiling BMC-H adopted a second organization (*Ass-B*) could therefore point to a structural trap delaying shell closure. To verify this hypothesis, two approaches were envisioned. First, we evaluated the interaction energy profile after gradually displacing the relative lateral disposition of hexamers. Umbrella sampling all-atom MD ^26^ were performed on hexamer couples extracted from PduA^*Sent*^ (3NGK), CcmK1^*6803*^ (3BN4) or CcmK4^*7942*^ (4OX6) structures, taken as representative of *Ass-A, Ass-B*, and *Ass-C* organizations, respectively (Fig. 4). Preliminarily, we measured the potential of mean force (PMF) that results from pulling apart the two partners (Fig. 4A). Despite comparable to calculations on CcmK2^*6803* 27^, CcmK1^*6803*^ binding energy was very weak, about 2 to 4 times smaller than for CcmK4 or PduA, respectively. When the PMF was screened along the reaction coordinate that leads from Ass-A to Ass-B (further extended on both sides), while hexamers were constrained flat, the highest stability was attained around crystal dispositions, supporting a preference of PduA^*Sent*^ for *Ass-A*, or of CcmK4^*7942*^ for *Ass-B/C* arrangements (Fig. 4B). Although CcmK1^*6803*^ did not manifest a clear preference, globally these data supported that arrangements occurring in 2D-tiling crystals represent energy minima. Besides, shallow profiles fitted with the high plasticity of BMC-H interfaces, indicating that transitions between different assemblies, separated by small energetic barriers (few k_B_T), should be feasible.

**Figure 4.**
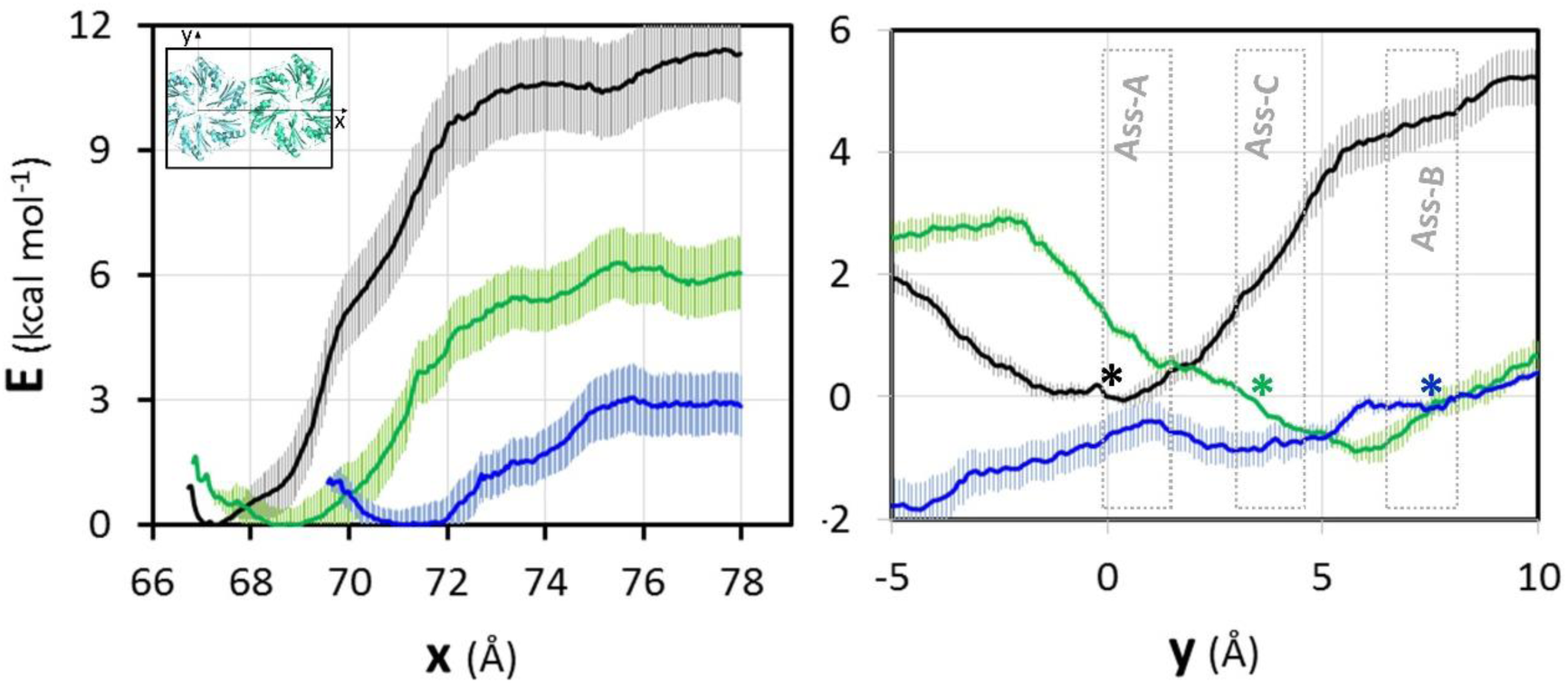
Potential of mean force (PMF) between two BMC-H hexamers. In the left panel, the PMF was calculated using umbrella sampling al atom MD simulations after separating progressively the two hexamers along the x-axis defined by the two center of masses (COM) of the hexamers in the starting X-ray disposition (inset). The origin of energy is taken at the minimum of the PMF so that the value of the energy at the largest distance provides an estimation of binding energies. For the right panel, the hexamers were gradually displaced along the orthogonal y-axis and the PMF was calculated using restraints to prevent bending, tilting and z-rotation Data for PduA^*sent*^ (3NGK) is plotted in black, in blue for CcmK4^6803^ (3BN4) and green for CcmK4^*7942*^ (40×6), including error bars estimated by bootstrapping. The approximate location of the different assembly modes is indicated in the right panel. The asterisks are to indicate the approximate disposition of the starting crystal for each case (following the mentioned color code).

### Behavior of reconfigured assemblies

The second approach consisted in launching MDs simulations on tri-hexamers of CcmK, EutM^*Ecol*^ and BMC-H^*Hoch*^ (remaining flat in crystal dispositions), after artificially repositioning each hexamer in *Ass-A* configuration. For that, each hexamer was superimposed individually on the different hexamers of the template PduA^*Sent*^ (3NGK) and potential clashes relaxed (see M&M). We decided to include RMM-H^*Msm*^ in the study, in view of its demonstrated potential to form nanotubes ^24^.

MD launched on reconfigured assemblies revealed significantly more stable than when starting from crystal dispositions, and collectively confirmed that *Ass-A* likely represents an arrangement competent for shell closure for most, if not all BMC-H (Fig. S14). Thus, BMC-H^*Hoch*^ and RMM-H^*Msm*^ behaved much like CsoS1A^*Hneap*^ or PduA^*Sent*^ in dynamics of Fig. 3. Also remarkable was CcmK4^*7942*^ curving trend, contrasting with the reproducible flatness of this protein when arranged as in the crystal. An exception was CcmK4^*6803*^, which remained flat. However, the inter-hexamer separation increased by almost 2 Å, similarly to CcmK2^*7942*^ (Table S4), something that could point to insufficiently relaxed starting structures. Since several bulky residues lie at the junction of the three CcmK4^*6803*^ hexamers and might hamper bending, we assessed a mutant with several residues replaced by corresponding residues from PduA^*Sent*^: R30N, Q53G, E54A, E85T and N86D. This mutant bent, albeit still less pronouncedly than other cases (Fig. S14).

### Experimental assembly of selected PduA mutants

Structural analysis (Fig. 1) and sequence alignments (Fig. S15A) indicate that ionic interactions mediated by residues corresponding to Arg28 and Asp49 of CcmK1^*6803*^ could contribute to hold some BMC-H in non-*Ass-A* organizations. The consequence of mutating the Asn29 of PduA^*Sent*^ into Arg was therefore evaluated. Similarly, the ionic bridge was disrupted in CcmK4^*7942*^ and BMC-H^*Hoch*^ *via* R29N or K28N mutations, respectively. In this chapter, we also investigated the effect of substituting Ala for the two key interfacial residues of PduA^*Sent*^: Lys26 and Arg79.

Overexpression of any of the three PduA^*Sent*^ mutants in BL21(DE3) caused a significant reduction of the proportion of elongated cells, when compared to the WT protein (Fig. S15B). The K26A mutation induced the strongest effect, when compared to R79A and N29R. On the contrary, disruption of the presumed ionic bridge led to a significant increase with both CcmK4^*7942*^ and BMC-H^*Hoch*^ mutants. SDS-PAGE analysis supported that all mutants continued to assemble inside cells, since most of protein was recovered after resolubilisation with 1M urea for all cases (Fig S15C).

Assembly of mutants recovered in UPF was then investigated by TEM. Only small patches were observed with K26A PduA^*Sent*^, structures that could not be unequivocally attributed to protein assemblies (Fig. S16A). The R79A mutant gave rise to compact 2D-patches (Fig. S16B). Structures resembling 40-80 nm amorphous nanotubes were seen, albeit rarely, together with a single sharp nanotube that could not be reproduced. With the N29R mutant, 2D-patches grew abundantly, sometimes accompanied by nanotube-like bundles (Fig. S16C). Differing from observations on the WT protein, hexamers were well defined inside the 2D-plateforms, appearing as strips arranged with approximately 120° angles, with long inter-hexamer separations (from 7.1 up to 9 nm, depending on direction), overall fitting to the occurrence of *Ass-B* arrangements with N29R PduA. Possibly due to higher compactness, hexamer definition decreased at platform edges, in nanotube bundles too. Also differing from WT, individual nanotubes were seen only once, and mixtures of motifs seem to coexist within the bundles. These data might point to assembly transitions that would be favored by subunit exchange and could trigger for instance wrinkling of 2D-patches towards nanotube-like bundles. On the other hand, 2D patches imaged for the R29N CcmK4^*7942*^ or K28N BMC-H^*Hoch*^, appeared more compact than those characterized by AFM for the respective WT proteins (Fig. S16D,E). However, nanotubes were not detected with any of the two mutants.

Intrigued by the profiles of cellular elongation, we decided to monitor assembly formation inside cells (Fig. 5, see also Fig. S17). Overexpression of WT PduA^*Sent*^ resulted in patterns compatible with 22-24 nm-wide nanotubes, although transverse sections failed to reveal described honeycomb motifs ^10, 15^. The K26A and R79A mutants gave rise to very similar compact assemblies, which almost filled up the cytosol. Stripped motifs, which were more evident with R79A PduA, were spaced by about 7 to 9 Å, suggesting the presence of piled sheets, as proposed before ^28^. Most interesting, the R29N PduA^*Sent*^ mutant did not form nanotubes (Fig. 5D and S17D). Instead, fingerprint-like arrangements, possibly single layered, were detected.

**Figure 5.**
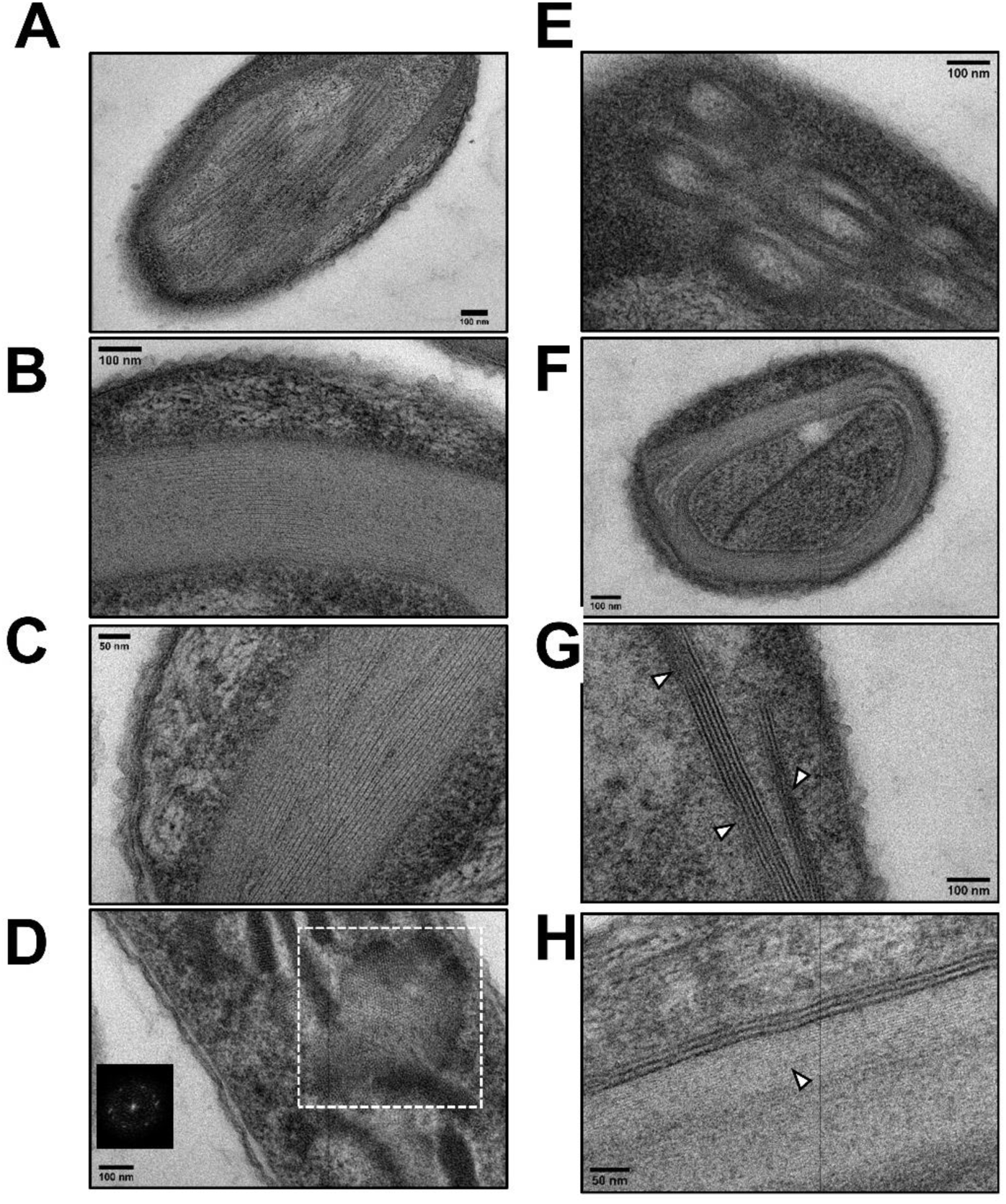
Single point mutant BMC-H Assemblies inside *E. coli*. Representative TEM views of cells overexpressing the next proteins: *A*, wild-type PduA^*Sent*^; *B*, K26A PduA^*Sent*^; *C*, R79A PduA^*Sent*^; *D*, N29R PduA^*Sent*^; *E*, wild-type BMC-H^*Hoch*^; *F*, K28N BMC-H^*Hoch*^ and *G-H*, R29N CcmK4^*7942*^. Stripped motifs occurred in cells overexpressing wild-type PduA, with 23-25 nm periodicities matching to typical nanotube dimensions described for this protein. No noticeable difference was seen between longitudinal or transverse views of cells. Extended 2D-patches imaged with K26A and R79A PduA mutants were identical, although striped motifs were more frequent with the R79A mutant. Motif separations of 6 to 9 Å patterns could point to transversal views of piled 2D layers, although laterally packed 1D-organized hexamers cannot be ruled out. Overexpression of the R29N PduA mutant (*D*) resulted in cells covered by 2D-assemblies with 6.8-8,1 Å periodicities (FFT treatment of outlined region shown as inset). *E*, BMC-H^*Hoch*^ revealed arrangements that fit to described Swiss-rolls. Motifs inside seem to repeat every 5.6 to 7 Å. The same periodicities were noticed for the K28N mutant (*F*). Assemblies looked more massive and compact than with the WT BMC-H^*Hoch*^. The rolled layers were absent from transverse views. *G-H*, The R29N CcmK4^*7942*^ resulted in big assemblies, comparable to those observed for PduA mutants. However, 12 to 16 Å-distant striped motifs reminiscent of nanotubes were noticed inside a majority of cells. These motifs were occasionally apposed on edges of more compact accompanying arrangements (triangles), suggesting assembly transitions. These and other high resolution images are available upon request for more detailed analysis (see also Fig. S17).

Intriguingly, most of cells displayed zebra-like patterns, something that might arise from 2D-layer wrinkling. While reported rolled layers were reproduced with BMC-H^*Hoch* 22, 29^ (Fig. 5E) and the structures were affected by replacement of Lys28 by Asn (Fig. 5F, S17F), the mutation did not cause appearance of nanotubes. However, structures reminiscent of nanotubes or nanowires accompanied other compact patches in a majority of cells overexpressing the R29N CcmK4^*7942*^ mutant (Fig. 5G-H and S17G).

## Discussion

Understanding how macromolecular structures as complex as BMC or BMC shells form is challenging. Implying the controlled assembly of hundreds to thousands of oligomeric subunits ^30^ deriving from several components, such processes are thought to rely in cooperativity phenomena. Relatively weak interactions, in the low *k*_B_T regime, normally govern contacts between pairs of shell components ^27^. That explains why coarse-grained models are precious tools to investigate parameters governing these processes ^12, 13^, in spite of being unsuitable for predictive work aiming to engineer new assemblies.

With the intention to contribute to this effort, this study sought to investigate global relationships between BMC-H experimental assembly behavior and predictions from all-atom MD simulations. Compiled experimental data pointed to a segregation of BMC-H in two major groups (Table S1), depending on trends to form flat assemblies (basically *β*-carboxysome CcmK) or rounded objects (e.g. Pdu or CsoS1 proteins). Within the first group could be included cases like BMC-H^*Hoch*^, found here or before to assemble flat *in vitro* ^22, 23^, and leading to rosette structures inside *E. coli* that are assumed to derive from spooling of quasi-flat carpets^22, 29, 31^. Nanotubes/spheroids and planar structures are not necessarily mutually exclusive, as demonstrate for example data presented here for PduA or PduJ (e.g. Fig. S4G-I), or before for several EutM homologs assembled at variable temperatures ^32^. Transitions between flat and curved structures might be one of the consequences of the relatively shallow interaction energy profiles calculated here for several BMC-H.

A notable discovery was that BMC-H experimental assembly behavior appears to be related to the type of organization found in crystals that exhibit internal 2D-layered organizations. Bending cases like PduA^*Sent*^, PduJ^*Sent*^ or CsoS1A^*Hneap*^ exhibited *Ass-A* dispositions, whilst other arrangements are noticed in structures of CcmK, EutM^*Ecol*^, EutM^*Cdif*^ or BMC-H^*Hoch*^. Most often, assembly units were laterally displaced by approximately 8-10 Å (*Ass-B*), when compared to *Ass-A*. If such relationship was correct, crystal data would indicate that proteins like CsoS1C^*Hneap*^, CsoS1^Pmar^ and BMC-H^*Ahyd*^ should form rounded structures, temptingly also BMC-H that attained *Ass-A* states even when mutated: CmcB^*Ecol*^ (7MN4, K25A-E55G mutant), CmcC^*Ecol*^ (7MPX, K25A-E35G) or CutR^*Sint*^ (6XPI, K66A). However, this rule is not obeyed by all BMC-H, since EutM is prone to form rounded structures but adopts *Ass-B* configurations in crystals. EutM complex behavior is illustrated by the varied types of structures formed depending on the homolog selected ^32^.

That hexamers could organize differently within 2D-layered crystals was already known since the early report of CsoS1A^*Hneap*^ crystal structure ^33^. At that time, side-to-side hexamer packing differences between CsoS1A (66.4 Å inter-hexamer separation) and CcmK structures (69.7 or 70.5 Å for CcmK1^*6803*^ or CcmK4^*6803*^, respectively) was argued to reflect means to attain compatibility among shell constituents or to regulate the porosity of the shell. The availability of many more structures nowadays rules out that differences were coincidental or induced by crystallization conditions, which spread considerably by pH (4.2 to 9.5) or additives within each group (Table S2), and occasionally overlapped between the two classes (compare for instance conditions with 4OX8 vs 4OX6). Indeed, PMF profiles estimated here supported that highest stabilization is attained in different flat dispositions of PduA^*Sent*^ and CcmK4^*7942*^ hexamers. Although CcmK1^*6803*^ profile was unexpectedly shallow, that CcmK never reached an *Ass-A* disposition in crystals (10 PDB entries) solidly points to global minima different from *Ass-A*. Yet, the structures of all reconstituted BMC shells prove that *Ass-A* is the mode of organization that conducts to closed assemblies (Table S2), also for shells composed of CcmK proteins ^16^.

All-atom MDs robustly demonstrated the experimental curving capabilities of *Ass-A*-organized BMC-H. Not only PduA^*Sent*^, but also all other *Ass-A* BMC-H rapidly and robustly bent. The known bending orientation was reproduced, i.e. the convex face lying towards the inner side of the compartment. In fact, structures averaged over the MD were strikingly similar to those found in minimalist shells, which all appear to derive from *Ass-A* arrangements. Comparison with data published for PduA^*Sent*^ bi-hexamers ^14^ indicate that assembly might be better simulated on tri-hexamer ensembles, although in our hands a preference for curved dispositions was even demonstrated on bi-hexamers extracted from minimalist BMC-H^*Hoch*^ and CcmK^*7418*^ shells. BMC-H with *Ass-B, Ass-C* or *Ass-D* organizations behaved less clear-cut, bending and even tilting values being much dispersed over the MD trajectories. Globally, however, *Ass-B* or *Ass-C* tri-hexamers seem to fluctuate around flat dispositions. This was seen for individual MD runs [e.g. CcmK4^*7942*^ (4OX6)], or was concluded from independent MDs [e.g. CcmK2^*6803*^ behavior in assemblies prepared from crystals 2A1B and 3CIM]. However, curving capabilities could be also demonstrated for these BMC-H when reconfigured in *Ass-A* dispositions, suggesting that transitions between assembly modes might be a means to control BMC closure.

The ionic pair between Arg/Lys at position 28 and Asp49 (CcmK1^*6803*^) likely contributes to maintain some BMC-H in *Ass-B* arrangements. This is supported by hexamer interspacing in 2D-patches imaged by TEM for the purified N29R PduA^Sent^, or conversely by the compact patches formed by R29N CcmK4^*7942*^, possibly by K28N BMC-H^*Hoch*^ too. Similarly, 2D-patches were seen in cells overexpressing N29R PduA^Sent^, contrasting with nanotubes typically observed with WT PduA, whereas structures compatible with nanotubes/nanowires occurred in cells expressing the R29N CcmK4^*7942*^. However, the latter, and K28N BMC-H^*Hoch*^ continued to assemble flat when purified, while nanotube-like bundles happened with N29R PduA^Sent^, suggesting that the mutations increased the chances of observing the inversed arrangements, but did not suffice to fully alter BMC-H assembly behavior.

Also challenging were efforts to clarify the atomic determinants behind spontaneous curvature. Difficulties are illustrated by the striking failure to disrupt the assemblies when key residues are mutated. This is the case of K26A, N29R and R79A PduA^*Sent*^ mutants studied here or before ^15^, of K28A and R78A BMC-H^*Hoch* 22, 29^, or K25A, R28A, D49A and R80A CcmK1^*6803* 21^. Assemblies continued to grow, in spite of the fact that interactions between constituting subunits are weak. One of these residues, corresponding to K26 of PduA^*Sent*^, is conserved at the interface of all BMC-H. Its mutation into Ala or Asp in PduA^*Sent*^ led to *Ass-B* arrangements in crystals (Table S2), while flat patches were imaged in *E. coli* (Fig. 5B), further reinforcing the link between crystal arrangements and curving trend. Energetic calculations proved that this residue and R79 are key interactors (Fig. S10). These data might match with the important role for curvature proposed for R79 ^14^. In that study, the mutation of PduA R79 in a ΔPduJ strain was shown to hamper the formation of nanotubes or even of Pdu BMC. TEM data presented here confirms this effect *in vitro*. Yet, mutation of other positions was known to have similar consequences [e.g. for K26A PduA ^15^ or K25A PduJ ^17^], as also demonstrated here for K26A and even N29R PduA^*Sent*^ mutants, altogether inviting caution before concluding on cause/effect relationships. In our hands, R79A did not exert any significant effect on curvature *in silico*. Among 21 mutants tested, only the interfacial K26 (PduA) completely and reproducibly abolished bending, but the effect was weaker in other BMC-H. The prediction was supported by the neat drop of the proportion of abnormally elongated BL12(DE3) with the K26A mutant, as compared to WT PduA, and by the direct observation of 2D organizations and absence of nanotubes within these cells. Interestingly, Pdu compartments were not recovered from *S. enterica* when the K26 of chromosomal PduA or K25 of PduJ were mutated into alanine, whereas compartments still formed with the R79A mutant and the protein was even co-purified with the BMC ^34^. Piled 2D sheets were also imaged by TEM for a K26A mutant of a variant of PduA from *Cit. freundii* expressed in *E. coli* ^15^. However, in this last study the R79A mutation elicited similar results, likewise in our study. Accordingly, the two residues might be proposed to play a concerted role. K26 side-chain is often modeled in crystals with a stretched conformation, lying antiparallel with regard to the same residue of the counter-interacting hexamer. Asp22, also fully conserved, contributes to hold such conformation. A consequence is that the positively-charged groups of K26 and R79 are brought closer, something that might impose an extended conformation to R79 sidechain and facilitate its insertion in a small pocket of the facing hexamer in *Ass-A* dispositions. Alternatively, PduA curvature might be only directly defined by the electrostatics around the K26 ammonium group. The R79A would exert its effect indirectly, for instance if the mutant was incapable of reaching the required *Ass-A* disposition, similarly to the N29R mutant. These processes would be out of reach for short MD simulations. Irrespective of the mechanisms that trigger curvature, our observations have implications for the interpretation of BMC biogenesis. *β*-carboxysome formation was proven to be two-step process in *Syn. elongatus* PCC 7942, the presence of pre-organized cargo preceding shell assembly ^7^. A similar study on Pdu^*Sent*^ compartments pointed instead to a less synchronized mechanism, with coexistence of *cargo-first* and *shell-first* events ^11^. A similar model could operate with α-carboxysomes, as indicate the gathering of CsoS2 scaffolding protein to portion of shells, which would precede recruitment of RuBisCO and final BMC closure ^35^, or the multiple layers of RuBisCO attached to the inner surface of partial α-carboxysome shells ^8, 36^. In view of that, and considering that BMC-H are the most abundant shell components, *Ass-B* might be seen as a structural trap that would delay premature shell closure of some BMC such as *β*-carboxysomes. Transit towards *Ass-A* dispositions would be triggered by contacts with motifs on cargo/scaffold components or, alternatively, upon intervention of auxiliary proteins such as CcmO or CcmP. Indeed, BMC-T co-expression was required to induce the formation of *Halothece* sp. PCC 7418 or BMC-H^*Hoch*^ empty shells, both inside cells ^16, 37^ or *in vitro* ^38^, while BMC-T presence was dispensable for the formation of RMM-H^*Msm*^ nanotubes ^38^, minimalist α-carboxysomes ^39^ or *Klebsiella pneumoniae* GRM2 shells ^40^. In the last study, cmcC from GRM2 was proposed to be a component endowed with high spontaneous curvature, in agreement with the crystal *Ass-A* organization of the *E. coli* K25A cmcC homolog ^41^. Future studies will be required to address the possible implication of BMC-T in mediating BMC-H assembly transitions, also to investigate the physiological consequences of altering the natural biogenesis pathway of a given BMC.

## MATERIALS AND METHODS

### Cloning, Expression, and Protein Purification

DNA sequences permitting the expression of the different BMC-H studied here are given in the excel Table S5, including flanking regions. These segments were integrated in between *XbaI* and *BlpI* sites of a pET26b vector using standard cloning methods. Plasmids for studies of *Synechocystis sp*. PCC8603 based on pET15b were described before ^19, 21^. After transformation of BL21(DE3) cells, protein expression was carried out in ZYM-5052 auto-induction media (4 mL) overnight at either 25°C or 37°C, following a described protocol ^19^. Cells pellets collected by centrifugation (4000 g) were lysed with 0.5 mL BugBuster extraction reagent (Sigma), supplemented with PMSF (1 mM, final conc.). Small aliquots were withdrawn and prepared for SDS-PAGE analysis (denatured at 95°C), corresponding to cellular fractions (CF). Soluble fractions (SF) were prepared similarly from material remaining in supernatants after centrifugation at 21000 g, 4°C for 15 min (pellets were kept at 4°C for studies of protein disassembly). Purification was performed on Vivapure 8 96-well cobalt-chelate microplate columns, following provider (VivaScience) instructions. Elution was effected with 300 μL of a 20 mM NaPi/300 mM NaCl/300 mM imidazole (pH 8.0) solution. EDTA (5 mM final conc.) was added immediately after elution and purified fraction (PF) aliquots withdrawn and denatured for SDS-PAGE analysis. Material remaining in pellets were re-treated with BugBuster, sonicated for 20 sec at 4°C and centrifuged at 21000g. After discarding supernatants, pellets were supplemented with 600 μL of a 20 mM NaPi/300 mM NaCl/10 mM imidazole (pH 8.2) solution containing 1 M urea. After resuspension with vigorous vortexing, the suspension was shaken for 3-5 hours at 4°C, with two interspersed rounds of 20 sec sonication. After 21000 g, urea-solubilized fraction (USF) aliquots were prepared for SDS-PAGE as other fractions, and the material was purified as indicated above on microplate columns. In that manner urea-purified fractions (UPF) were prepared.

The remaining of material eluted from the purification columns, which permitted to prepare PF and UPF, was dialyzed at 4°C against 10 mM NaPi/150 mM NaCl (pH 6.5) using 3500 Da MWCO Pure-A-Lyzer devices. Two steps were carried out, theoretically permitting a > 10^4^ fold decrease of initial buffer and urea concentrations. When necessary, proteins were concentrated to 1-2 mg/mL using Vivaspin 500 centrifugal concentrators (PES, 10 kDa MWCO).

### Phase contrast microscopy

Cells growing in LB supplemented with the corresponding antibiotic were induced at exponential phase (OD^280nm^ = 0.6-0.8) with IPTG (200 μM). Growth was pursued at 37°C for 6 h before visualization under an oil immersed 100x objective lens with a DM4000B Leica microscope (DFC 450C caméra and EF6000 laser). Image analysis was carried out with ImageJ using an automated procedure that consisted in i) subtracting signal background, ii) setting-up a common threshold, and iii) rejecting small objects, well below the cell size.

### Atomic Force Microscopy imaging

AFM experiments were carried out exactly as previously described ^21^, after diluting protein samples in 10 mM NaPi, 300 mM NaCl pH 6.5 (*Buffer P*). When indicated, this solution was supplemented with 1 mM MgCl_2_ (final conc.) with the intention to inverse mica surface charge.

### Transmission electron microscopy

Specimens were prepared for electron microscopy using the conventional negative staining procedure. After diluting proteins in *Buffer P* to 0.05-0.15 mg/mL, 10 µL drops were deposited on parafilm for 20 min. After absorbing material on glow-discharged Formvar carbon-coated grids for 2 min, grids were blotted and negatively stained with uranyl acetate (1%) for 1 min.

For analysis of cell contents, induction was effected with IPTG as indicated above. After 6h, cells were pelleted and resuspended in 2,5% glutaraldehyde and 2% paraformaldehyde in cacodylate buffer (0.1 M, pH 7.2). After 2 h at RT, cells were post-fixed with 1% OsO_4_ in the same buffer. After pelleting and concentrating in agarose, cells were treated for 1 h with 2% aqueous uranyl acetate. The samples were then dehydrated in a graded acetone series and embedded in Epon. After 48 h of polymerization at 60 °C, ultrathin sections (80 nm thick) were mounted on 200 mesh Formvar-carbon-coated copper grids. Sections were stained with Uranyless and lead citrate.

A JEM-1400 electron microscope (JEOL Inc, Peabody, MA, USA) operating at 80 kV, equipped with a Gatan Orius digital camera (Gatan Inc, Pleasanton, CA, USA) was used for imaging at different magnifications.

### Cryoelectron microscopy (Cryo-EM)

3.5µL sample drops were deposited on parafilm for 20 min. Glow-discharged lacey carbon grids were then placed onto the drops and incubated for 1min. Grids were transferred to the thermostatic chamber of a Leica EM-GP automatic plunge freezer, set at 20°C and 95% humidity. Excess solution was removed by blotting with Whatman n°1 filter paper for 0.2-0.6 seconds, and the grids were immediately flash frozen in liquid ethane at −185°C. Frozen specimens were placed in a Gatan 626 cryo-holder, and cryo EM was carried out on a Jeol 2100 microscope, equipped with a LaB_6_ cathode and operating at 200 kV, under low dose conditions. Images were acquired with SerialEM software, with defocus of 1.5– 2.5 μm, on a Gatan US4000 CCD camera. This device was placed at the end of a GIF Quantum energy filter (Gatan, Inc.), operated in zero-energy-loss mode, with a slit width of 25 eV. Images were recorded at a nominal magnification of 4,000 corresponding to calibrated pixel sizes of 1.71Å.

### All-atom Molecular Dynamics Simulations

Assemblies composed of three hexamers were prepared from available crystallographic structures (PDB ID indicated in Table S2) after applying translation and symmetry operations. Glycerol and other crystallographic ligands were removed (sulfate ions associated to CsoS1A were deleted, or not, without evident difference). Ensembles were embedded in cuboid cells with dimensions extending 20 nm around all protein atoms, which were filled with explicit solvent. Each system was neutralized with NaCl (0.9 % final concentration). Periodic boundary conditions were applied and unless otherwise mentioned the YASARA AMBER14 force field was selected. The cut-off for the Lennard-Jones potential and the short range electrostatics was 8 Ǻ. Long-range electrostatics were calculated using the Particle Mesh Ewald (PME) method with a grid spacing <0.1 nm, 4th order PME-spline, and PME tolerance of 10^−5^ for the direct space sum. YASARA’s pKa utility was used to assign pKa values at pH 7.0. The entire system was energy-minimized using steepest descent minimization, followed by a simulated annealing minimization until convergence (<0.05 kJ/mol/200 steps). The equations of motions were integrated with a multiple time step of 2.5 fs for bonded interactions and 5.0 fs for non-bonded interactions at a temperature of 298K and a pressure of 1 atm (NPT ensemble). At least two simulations were launched on each case, with attribution of random initial atomic velocities (20 ns/run, unless otherwise indicated). Intermediate MD snapshots were recorded every 250 ps. Simulations were carried out in a 16-core CPU PC exploiting GPU capabilities (NVIDIA GeForce GTX 1080), and lasted typically 50 to 60 hours per 20 ns run.

For simulations of *Ass-A* reconstituted assemblies, hexamers extracted from target PDB structures were superimposed individually on the different hexamers of the template PduA^*Sent*^ (3NGK) tri-hexamer. For most cases, sterical clashes around R30, the D51-E54 segment and the R82-N86 region (CcmK4^*6803*^ numbering) were alleviated by adapting the side-chain conformation to reproduce those present in the corresponding residue of PduA (3NGK). Special attention was given to the CcmK conserved Arg30. Its side-chain conformation was adapted to reproduce the orientation observed for Arg28 in the cryo-EM structure of the CcmK^*7418*^ shell (6OWF). In CcmK4^*6803*^, the Arg30, Gln53 and Glu54 collapse close to the C3 axes of symmetry of the tri-hexamer. These side-chains were therefore adapted manually.

Dihedral angle and distance analysis was performed with adapted Python scripts executed under Pymol (https://www.pymol.org/). Tilting was measured among main-chain Cα atom positions of M24 and Ile18 (PduA^*Sent*^, corresponding residues in other BMC-H) from two monomers of a given hexamer with regard to same symmetric residues of the interacting hexamer counterpart, as described^21^. Bending values were based on Cα atoms of S27 and Ile38 from one of the interfacial monomers and the same residues on the symmetric monomer of the opposite hexamer. For plane representations, structures averaged over the MD were first superimposed on the corresponding crystal structure. Only main-chain atoms from one of the hexamers was used for the superimposition. Next, each hexamer was represented by the plane containing V54 Cα atom (PduA^*Sent*^) of the three monomers of the given hexamer that compose the interface. In other representations, each hexamer was represented by its center of mass, calculated considering only main-chain atoms of core residues (res 1 to 90) from the six monomers. Structural figures were also prepared with Pymol.

Data from MD trajectories snapshots, either in YASARA .sim format or as pdb files, as well as AFM or TEM data presented in this study are available upon request.

### Umbrella Sampling Molecular Dynamics Simulations

Assemblies of two hexamers of PduA^*Sent*^ (3NGK), CcmK1^*6803*^ (3BN4), and CcmK4^*7942*^ (4OX6) were prepared at pH 7.0 from the available crystal structure using ProteinPrepare (https://playmolecule.com/proteinPrepare/). All atom MD simulations were performed using GROMACS (version 2021.1), with Amber ff99SB-ILDN force field. All other conditions were as mentioned above, with the difference that the cut-off for the Lennard-Jones potential and the short-range electrostatics was established at 10 Ǻ. The entire system was energy-minimized using steepest descent minimization, followed by a constant volume, constant temperature (NVT) equilibration at T=298 K for 100 ps with the backbone of both hexamers restrained with a force constant of 1000 kJ/mol/nm^2^.

Umbrella sampling simulations were performed with translation and rotation of the first hexamer prevented with harmonic restraints (force constant of 1000 kJ/mol/nm^2^) applied to six C_α_ atoms (one per monomer, near its COM). Binding energies were evaluated using the distance between the two hexamers COM as reaction coordinate. A steered simulation pulling on the second hexamer with a force constant of 1000 kJ/mol/nm^2^ at a rate of 10 nm/ns was used to generate configurations at 1 Å steps along the reaction coordinate. A total of 15 windows per case were therefore processed through 2 ns of NPT equilibration at 1 atm and 10 ns of production simulations to finally calculate the PMF between the two hexamers using the Weighted Histogram Analysis Method (WHAM) ^26^. The error was estimated using bootstrapping. The PMF as a function of the hexamer lateral displacement as reaction coordinate was calculated similarly, with the difference that the second hexamer was progressively displaced using UCFS Chimera from the crystal disposition, at 1 Å steps in a range between −15 Å and + 15 Å. In those simulations the bending, tilting, and z-rotation angles were restrained to zero with harmonic potentials with a force constant 1000 kJ/mol/rad^2^ to keep the two hexamers planar.

## SUPPORTING INFORMATION

Next files are provided as supplementary data: **Table S1** (DOC). Summary of nano-assemblies characterized here or previously for individual BMC-H; **Table S2** (XLSX). Analysis of hexamer dispositions in crystal structures with piles of tiling BMC-H; **Table S3** (DOC). Analysis of structural changes of tri-hexamers BMC-H assemblies occurring during MDs trajectories; **Table S4** (DOC). Structural changes occurring during MDs when BMC-H tri-hexamers are reconfigured as in *Ass-A* organizations; **Table S5** (XLSX). DNA sequences coding for BMC-H characterized experimentally in this work. And next figures (all PDF): **Figure S1**. Impact of overexpression of BMC-H on *E. coli* size and recovery of potential assemblies via urea solubilization; **Figure S2 (PDF)**. Potential CcmK assembly inside *E. coli* studied by TEM; **Figure S3**. BMC-H assembly visualized by AFM; **Figure S4**. Analysis by TEM of BMC-H assembly; **Figure S5**. Characterization of PduA, PduJ and CsoS1A assemblies by cryo-EM; **Figure S6**. Plot of tilting and bending angles measured in snapshots collected in the course of MD simulations on BMC-H tri-hexamer ensembles; **Figure S7**; Graphic view of results of MD simulations on bi-hexamers extracted from BMC shells; **Figure S8**. Monitoring RMSD evolution of PduA residues over MD simulations; **Figure S9**. Sidechain movements of selected PduA residues during MD simulations; **Figure S10**. Energetic contribution of PduA residues to the stabilization of the bent assembly; **Figure S11**. Impact on MD behavior of the PduA tri-hexamer when individual residues are mutated into alanine; **Figure S12**. Consequences for PduA assembly behavior of the replacement of Lys26 by other residue types; **Figure S13**. MD behavior of *Ass-A* tri-hexamer BMC-H assemblies with interfacial Lys and Arg residues mutated into alanine; **Figure S14**. Dynamic behavior of planar-behaving tri-hexamers when reconfigured in *Ass-A* disposition; **Figure S15**. Impact of overexpression of BMC-H with selected interfacial residue mutations on *E. coli* size and on the recovery of potential assemblies via urea solubilization; **Figure S16**. Alteration of BMC-H assembly by single point mutations of interfacial residues; **Figure S17**. Characterization of assemblies formed by BMC-H single point mutants inside *E. coli*.

## AUTHOR CONTRIBUTIONS

L.F.G.-A.: Designed study, performed experiments, MD simulations, analyzed data and wrote the manuscript; V.S. and S.B.: collected electron microscopy images; I.C. and E.L. AFM acquisition; S.C.-C. phase-contrast microscopy; M.F.-C. performed MDs with CHARMM; G.T. discussions and revised manuscript; D.R. evaluation of PMF energy profiles, discussions and participation to manuscript writing.

## NOTES

The authors declare no conflict of interest

