## Supplemental Table S1 for "Inferring assembly-curving trends of bacterial micro-compartment shell hexamers from crystal structure arrangements"

**Table S1. Nano-assemblies characterized for individual BMC-H.** Data were compiled from references indicated in the last column. Objects were imaged by TEM directly after protein expression inside living cells (generally *E. coli*), or by TEM, cryo-EM or AFM with purified proteins (*in vitro*). Curved-implying objects are highlighted with blue letters, black for flat structures. NA: not applicable.

| **BMC-H** | **Species** | **Tech.** | **Assemblies**  **in cells** | **Assemblies**  ***in vitro*** | **Details** | **Ref.** |
| --- | --- | --- | --- | --- | --- | --- |
| **CcmK1** | Hal. sp. Pcc7418 | TEM | None visible | Not done |  | 1 |
| **CcmK1** | Syn. sp. PCC 6803 | AFM | Not applicable (NA) | **2D assemblies** (cup-like patches) |  | 2 |
| **CcmK1** | Syn. sp. PCC 6803 | AFM | NA | **2D assemblies** (cup-like patches) |  | This study |
| **CcmK1** | Syn. sp. PCC 6803 | TEM | Bad defined motifs | **2D assemblies** (cup-like patches) |  | This study |
| **CcmK1** | Syn. sp. PCC 6803 | TEM | Not done | **2D-crystal assemblies** | lipid-air interface | 3 |
| **CcmK2** | Hal. sp. Pcc7418 | TEM | None visible | Not done |  | 1 |
| **CcmK2** | Syn. elongatus PCC 7942 | AFM | Not done | **2D assemblies** (occasionally with stripes) |  | 2 |
| **CcmK2** | Syn. elongatus PCC 7942 | AFM | NA | **2D flat assemblies** |  | This study |
| **CcmK2** | Syn. elongatus PCC 7942 | TEM | None visible | Not done |  | 4 |
| **CcmK2** | Syn. elongatus PCC 7942 | TEM | None visible | None visible |  | This study |
| **CcmK2** | Syn. sp. PCC 6803 | AFM | NA | **2D assemblies** (occasionally with stripes) |  | 2 |
| **CcmK2** | Syn. sp. PCC 6803 | AFM | NA | **2D assemblies** (occasionally with stripes) |  | This study |
| **CcmK2** | Syn. sp. PCC 6803 | TEM | Not mentioned | **Spheroids (30-50 nm)** |  | 5 |
| **CcmK2** | Syn. sp. PCC 6803 | TEM | None visible | No assemblies |  | This study |
| **CcmK2** | Syn. sp. PCC 6803 | TEM | Not done | **2D-crystal assemblies** (stacks) | lipid-air interface | 3 |
| **CcmK2** | T. elongatus BP-1 | TEM |  | **bodies** (100-300 nm) | Not clear, artifacts? | 6 |
| **CcmK3** | Syn. sp. PCC 6803 | TEM | Inclussion bodies No visible assemblies | Not possible |  | This study |
| **CcmK4** | Syn. elongatus PCC 7942 | TEM | None visible | Not done |  | 4 |
| **CcmK4** | Syn. elongatus PCC 7942 | AFM | NA | **2D flat assemblies** |  | This study |
| **CcmK4** | Syn. elongatus PCC 7942 | TEM | None visible | None visible |  | This study |
| **CcmK4 R29N** | Syn. elongatus PCC 7942 | TEM | **Filaments** &  **piled 2D layers** | **2D assemblies** |  | This study |
| **CcmK4** | Syn. sp. PCC 6803 | AFM | NA | **2D assemblies** |  | 2 |
| **CcmK4** | Syn. sp. PCC 6803 | AFM | NA | **2D assemblies** |  | This study |
| **CcmK4** | Syn. sp. PCC 6803 | TEM | None visible | No assemblies |  | This study |
| **CcmK4** | Syn. sp. PCC 6803 | TEM | Not done | **2D-crystal assemblies** (stacks) | lipid-air interface | 3 |
| **BMC-H** | Hal. ochraceum | TEM | **Swiss-rolls** |  |  | 7 |
| **BMC-H** | Hal. ochraceum | AFM | NA | **2D flat patches** | Untagged  presence of Mg2+ | 7 |
| **BMC-H** | Hal. ochraceum | AFM | NA | None visible |  | This study |
| **BMC-H** | Hal. ochraceum | TEM | None visible | **2D flat assemblies** |  | 8 |
| **BMC-H** | Hal. ochraceum | TEM | **Swiss-rolls** | Not done |  | 4 |
| **BMC-H** | Hal. ochraceum | TEM | **Swiss-rolls** | Not done |  | 1 |
| **BMC-H** | Hal. ochraceum | TEM | **Swiss-rolls** | **2D flat assemblies** |  | This study |
| **BMC-H** | Hal. ochraceum | TEM | **Swiss-rolls** |  |  | 9 |
| **BMC-H K28A** | Hal. ochraceum | AFM | NA | **2D flat patches**  two-stacked and larger than WT | slower exchange dynamics than WT | 7 |
| **BMC-H K28A** | Hal. ochraceum | TEM | Amorphous open rolls? | Not done |  | 1 |
| **BMC-H K28P** | Hal. ochraceum | TEM | **Nanotubes**  or layered **2D-sheets**? | Not done |  | 1 |
| **BMC-H K28N** | Hal. ochraceum | TEM | Piled **2D assemblies**? | **2D flat assemblies** |  | This study |
| **BMC-H R78A** | Hal. ochraceum | AFM | NA | **2D flat patches** smaller patches than WT | exchange dynamics  comparable to WT | 7 |
| **BMC-H R78A** | Hal. ochraceum | TEM | Incipient rolls? | Not done |  | 1 |
| **BMC-H** | Hal. ochraceum | X-ray | NA | **icosahedral shell** | artificial fusion of two BMC-H | 10 |
| **CsoS1A** | Hal. neapolitanus | AFM | NA | **spheroids** |  | This study |
| **CsoS1A** | Hal. neapolitanus | TEM | Not done | **Nanowires** |  | This study |
| **CsoS1A** | Hal. neapolitanus | c-EM | NA | **nanotubes, spheroids** |  | This study |
| **EtuA** | Clos.kluyveri | TEM | **Nanotubes** | Not soluble using urea |  | 11 |
| **EutM** | Clos. difficile F. gelatini T. saccharolyt. | TEM | Not mentioned | **Fibres and 2D-assemblies** |  | 12 |
| **EutM** | Clos. difficile | TEM | amorphous **swiss-rolls** or irregular **nanotubes?** |  |  | 13 |
| **EutM** | Escherichia coli | AFM | NA | **Spheroids** (rarely) |  | This study |
| **EutM** | Escherichia coli | TEM | Not done | **nanowires** (rarely) |  | This study |
| **EutM** | Sal. enterica | TEM | Thick **filaments** | Not mentioned |  | 14 |
| **EutM** | S. enterica M. hydrocarb. T. linaloolentis C. thermarum D. thermocist. P. hadalis A. metalliredigens F. gelatini | TEM | Not mentioned | rolled-up wide **nanotubes** not completely closed |  | 12 |
| **EutS** | Clos. difficile |  | None visible |  |  | 13 |
| **EutS** | Sal. enterica | TEM | **spheroid/polyhedra?** | Not mentioned |  | 14 |
| **PduA** | Cit. freundii | TEM | **Nanotube** bundles | Not mentioned |  | 15 |
| **PduA** | Cit. freundii | TEM | **Nanotubes** | Not done |  | 1 |
| **PduA** | Sal. enterica | AFM | NA | **spheroids** and/or **nanotubes?** |  | This study |
| **PduA** | Sal. enterica | TEM | **Rod-like** objects | Not mentioned | Inclusion bodies  solubilized 6M Gndn:HCl | 16 |
| **PduA** | Sal. enterica | TEM | Not done | **Nanotubes, nanowires  2D flat assemblies** |  | This study |
| **PduA** | Sal. enterica | c-EM | NA | **Nanotubes** |  | This study |
| **PduA** | Sal. enterica | TEM | **Nanotubes** | Not done |  | 4 |
| **PduA** | Cit. freundii | TEM | Not mentioned | **Nanotubes (20 nm)** |  | 17 |
| **PduA** | Sal. enterica | TEM | **Nanotube** bundles | Not mentioned |  | 18 |
| **PduA*** | Cit. freundii | TEM | **Filaments/nanotubes** | Not mentioned | Artificial extension | 15 |
| **PduA*** | Cit. freundii | TEM | **Nanotube** bundles | Not mentioned | Artificial extension | 19 |
| **PduA*** | Cit. freundii | TEM | **Nanotube** bundles | Not mentioned | C-ter modified | 20 |
| **PduA’** | Sal. enterica | X-ray | NA | **Dodecahedral cage** | Circular permutation Organized as pentamer | 21 |
| **PduA K26A** | Sal. enterica | TEM | None visible | Not mentioned |  | 18 |
| **PduA K26A** | Sal. enterica | TEM | layered **2D sheets** | not clear |  | This study |
| **PduA* K26A** | Cit. freundii | TEM | layered **2D sheets** | Not mentioned | Artificial extension | 19 |
| **PduA* K26D** | Cit. freundii | TEM | None visible | Not mentioned | Artificial extension | 19 |
| **PduA R79A** | Sal. enterica | TEM | piled **2D sheets** | **2D assemblies** |  | This study |
| **PduA* R79A** | Cit. freundii | TEM | layered **2D sheets** | Not mentioned | Artificial extension | 19 |
| **PduA* V51A** | Cit. freundii | TEM | **Nanotube** bundles | Not mentioned | Artificial extension | 19 |
| **PduA* V51D** | Cit. freundii | TEM | None visible | Not mentioned | Artificial extension | 19 |
| **PduA N29R** | Sal. enterica | TEM | **2D assemblies** | **2D assemblies** and **wrinckled patches** |  | This study |
| **PduJ** | Sal. enterica | TEM | **Nanotube** bundles | Not mentioned |  | 18 |
| **PduJ K25A** | Sal. enterica | TEM | None visible | Not mentioned |  | 18 |
| **PduJ** | Sal. enterica | AFM | NA | small **2D flat assemblies** |  | This study |
| **PduJ** | Sal. enterica | TEM | Not done | **Nanotubes  2D flat assemblies** |  | This study |
| **PduJ** | Sal. enterica | c-EM | NA | **Nanotubes  2D flat assemblies** |  | This study |
| **RmmH** | Myc. smegmatis | AFM | NA | **Spheroids?** |  | This study |
| **RmmH** | Myc. smegmatis | TEM | **Nanotubes** | **nanotubes** | Triton X100 purified 1 to 9 mg/mL Disassembly < 1 mg/mL | 22 |
| **RmmH** | Myc. smegmatis | TEM | Not done | Amorphous fibers? | Lower concentrations than previous | This study |
| **RmmH** | Myc. smegmatis | TEM | **Nanotubes** | Not done |  | 4 |
| **RmmH** | Myc. smegmatis | TEM | **Nanotubes** | Not done |  | 1 |
| **RmmH** | Myc. smegmatis | TEM | **Nanotubes** | **Nanotubes** | cleavage of  SUMOylated construct | 9 |
