## Supplemental Table S3 for "Inferring assembly-curving trends of bacterial micro-compartment shell hexamers from crystal structure arrangements"

**Table S3 – Structural changes of tri-hexamers assemblies occurring during MDs trajectories.**

| **Protein** | **PDBid** | **MD #** | **Tilting** | | | **Bending** | | | **Distance** | |
| --- | --- | --- | --- | --- | --- | --- | --- | --- | --- | --- |
|  |  |  | **Deg** | **(+/-)** | **Aver** | **Deg** | **(+/-)** | **Aver** | **Å** | **Aver** |
| **CsoS1A*^Hneap^*** | **2G13** | **1** | -3,9 | 2,4 | **-3,8** | -22,7 | 4,6 | **-23,1** | -0,6 | **-0,7** |
|  |  | **2** | -3,5 | 2,7 |  | -23,5 | 4,5 |  | -0,7 |  |
|  |  | **3** | -4,6 | 2,9 |  | -25,7 | 7,3 |  | -1,1 |  |
|  |  | **4** | -3,3 | 3,6 |  | -20,6 | 6,4 |  | -0,5 |  |
| **CsoS1A*^Hneap^*** | **2EWH** | **1** | -2,7 | 2,9 | **-2,6** | -16,2 | 4,9 | **-20,8** | 0,0 | **-0,5** |
|  |  | **2** | -3,1 | 2,8 |  | -23,3 | 5,0 |  | -0,8 |  |
|  |  | **3** | -2,1 | 3,7 |  | -22,9 | 5,7 |  | -0,6 |  |
| **CsoS1C*^Hneap^*** | **3H8Y** | **1** | -3,5 | 4,9 | **-4,1** | -27,1 | 7,3 | **-29,4** | -1,1 | **-1,6** |
|  |  | **2** | -4,2 | 3,3 |  | -29,2 | 4,7 |  | -1,5 |  |
|  |  | **3** | -4,6 | 2,6 |  | -31,8 | 5,0 |  | -2,0 |  |
| **PduA*^Sent^*** | **3NGK** | **1** | -2,2 | 3,0 | **-2,4** | -26,6 | 5,4 | **-26,6** | -1,2 | **-1,0** |
|  |  | **2** | -2,6 | 3,2 |  | -26,6 | 6,1 |  | -0,9 |  |
| **PduJ*^Sent^*** | **5D6V** | **1** | -1,4 | 6,9 | **2,0** | -2,4 | 7,9 | **-15,8** | 0,9 | **0,1** |
|  |  | **2** | 3,0 | 4,2 |  | -24,8 | 7,8 |  | -0,4 |  |
|  |  | **3** | 4,4 | 6,4 |  | -20,0 | 10,7 |  | -0,1 |  |
| **BMC-H*^Ahyd^*** | **4QIV** | **1** | 1,9 | 7,7 | **2,5** | -26,1 | 4,3 | **-26,3** | -2,3 | **-1,1** |
|  |  | **2** | 3,1 | 3,1 |  | -26,5 | 4,5 |  | 0,1 |  |
| **CcmK1*^6803^*** | **3BN4** | **1** | 3,0 | 7,5 | **1,3** | -1,3 | 10,4 | **0,5** | 0,8 | **1,2** |
|  |  | **2** | -0,5 | 10,6 |  | 2,3 | 11,2 |  | 1,5 |  |
| **CcmK1*^6803^*** | **3DN9** | **1** | -1,1 | 7,3 | **-0,9** | 17,0 | 14,6 | **10,4** | 1,6 | **1,3** |
|  |  | **2** | -0,7 | 6,9 |  | 3,9 | 9,5 |  | 1,1 |  |
| **CcmK2*^6803^*** | **2A1B** | **1** | -6,1 | 15,8 | **-10,1** | 16,3 | 18,4 | **10,5** | 0,7 | **2,9** |
|  |  | **2** | -14,2 | 19,5 |  | 4,8 | 20,2 |  | 5,1 |  |
| **CcmK2*^6803^*** | **3CIM** | **1** | -3,1 | 16,1 | **-4,2** | -2,7 | 19,0 | **-14,4** | 1,2 | **-0,7** |
|  |  | **2** | -5,3 | 8,9 |  | -26,0 | 12,2 |  | -2,7 |  |
| **CcmK4*^6803^*** | **6SCR** | **1** | -6,8 | 15,8 | **1,8** | 1,1 | 18,1 | **2,0** | 0,6 | **0,5** |
|  |  | **2** | 10,4 | 16,1 |  | 2,8 | 15,6 |  | 0,4 |  |
| **EutM*^Ecol^*** | **3MPW** | **1** | 0,3 | 4,8 | **-0,1** | -4,4 | 10,4 | **-3,6** | 0,1 | **-0,2** |
|  |  | **2** | -0,3 | 13,7 |  | -30,4 | 16,1 |  | -2,8 |  |
|  |  | **3** | -4,0 | 11,0 |  | 4,2 | 14,7 |  | 0,2 |  |
|  |  | **4** | 3,8 | 7,3 |  | 16,4 | 11,6 |  | 1,8 |  |
| **EutM*^Ecol^*** | **3MPY** | **1** | -1,9 | 7,0 | **-1,4** | 12,0 | 11,4 | **2,5** | 0,8 | **0,6** |
|  |  | **2** | -0,9 | 9,7 |  | -6,9 | 11,4 |  | 0,3 |  |
| **BMC-H*^Hoch^*** | **5DJB** | **1** | 2,3 | 6,5 | **-0,4** | -5,3 | 9,1 | **-9,9** | 0,9 | **0,4** |
|  |  | **2** | -1,1 | 6,9 |  | -11,6 | 9,4 |  | 0,9 |  |
|  |  | **3** | -2,5 | 9,2 |  | -12,7 | 11,0 |  | -0,5 |  |
| **CcmK2*^6803^*** | **3DNC** | **1** | -2,6 | 6,0 | **-0,3** | -37,5 | 8,4 | **-28,1** | -1,8 | **-1,4** |
|  |  | **2** | 2,0 | 9,6 |  | -18,7 | 14,7 |  | -1,1 |  |
| **CcmK4*^7942^*** | **4OX6** | **1** | -0,9 | 2,5 | **-1,0** | -7,4 | 4,3 | **-7,0** | 1,0 | **1,2** |
|  |  | **2** | -1,2 | 3,4 |  | -6,7 | 3,6 |  | 1,3 |  |
| **CcmK2*^7942^*** | **4OX7** | **1** | -2,6 | 14,6 | **-0,5** | -7,3 | 23,8 | **-11,6** | -1,5 | **-1,4** |
|  |  | **2** | 1,6 | 18,9 |  | -15,9 | 19,2 |  | -1,3 |  |
| **EutL*^Ecol^***  **(BMC-T)** | **3I87** | **1** | -1,4 | 4,5 | **-0,6** | 1,8 | 7,1 | **2,2** | 0,8 | **0,5** |
|  |  | **2** | 0,2 | 5,1 |  | 2,6 | 6,7 |  | 0,2 |  |

Tilting and bending values correspond to the average of deviations measured between each MD snapshot structure and the crystal structure. The averages combine the three measurements between each couple of hexamers in the tri-hexamer assembly. For bending angles, negative sign indicates orientation towards BMC-H convex side. In occasions, local structural distortions around residues selected for calculation of angles could result in misleading values. Angles therefore need to be contrasted with more faithful plane representations prepared taking the center of mass (COM) of hexamers (see Fig. 3). Deviation of distances were calculated taking the coordinates of the COM of each hexamer in the MD average structure with regard to the crystal. Only main-chain atom coordinates of residues 1-90 in each monomer were considered for the estimation of each hexamer COM. The value is the average of the three inter-hexamer measurements. The Aver column provides an average of all independent MD runs. For PduJ*^Sent^* (5D6V), the alanine mutated residue in position 26 was replaced by the lysine residue of the wild-type protein.
