## Supplemental Table S4 for "Inferring assembly-curving trends of bacterial micro-compartment shell hexamers from crystal structure arrangements"

**Table S4 – Structural changes occurring during MDs trajectories when BMC-H tri-hexamers are reconstituted in *Ass-A* mode.**

| **Protein** | **PDBid** | **MD #** | **Tilting** | | | **Bending** | | | **Distance** | |
| --- | --- | --- | --- | --- | --- | --- | --- | --- | --- | --- |
|  |  |  | **Deg** | **(+/-)** | **Aver** | **Deg** | **(+/-)** | **Aver** | **Å** | **Aver** |
| **CcmK1*^6803^*** | **3BN4** | **1** | 1,0 | 4,8 | **0,0** | -21,9 | 9,2 | **-21,1** | -1,1 | **-1.1** |
|  |  | **2** | -0,9 | 3,6 |  | -20,3 | 6,3 |  | -1,1 |  |
| **CcmK2*^6803^*** | **2A1B** | **1** | 0,3 | 5,4 | **1,3** | -18,4 | 5,0 | **-21,2** | 0,5 | **0.3** |
|  |  | **2** | 2,3 | 3,8 |  | -24,1 | 6,6 |  | 0,2 |  |
| **CcmK2*^7942^*** | **4OX7** | **1** | 4,2 | 6,3 | **4,0** | -11,5 | 9,1 | **-11,9** | 1,9 | **1.8** |
|  |  | **2** | 3,8 | 6,1 |  | -12,4 | 7,0 |  | 1,7 |  |
| **CcmK4*^6803^*** | **6SCR** | **1** | 2,0 | 4,2 | **2,9** | -4,8 | 4,5 | **-2,4** | 1,8 | **1.9** |
|  |  | **2** | 3,8 | 4,5 |  | -0,0 | 3,6 |  | 1,9 |  |
| **CcmK4*^7942^*** | **4OX6** | **1** | -3,6 | 4,8 | **-2,2** | -21,1 | 5,4 | **-20,9** | -0.0 | **-0.1** |
|  |  | **2** | -0,8 | 2,8 |  | -20,8 | 7,0 |  | -0.2 |  |
| **EutM*^Ecol^*** | **3MPW** | **1** | 2,4 | 4,8 | **1,6** | -27,6 | 7,0 | **-23,6** | -0.2 | **0.1** |
|  |  | **2** | 0,9 | 9,4 |  | -19,7 | 9,1 |  | 0.4 |  |
| **BMC-H*^Hoch^*** | **5DJB** | **1** | -0,7 | 2,9 | **-0.2** | -28,8 | 7,5 | **-29,5** | -1.2 | **-1.2** |
|  |  | **2** | 0,4 | 2,9 |  | -30,2 | 6,1 |  | -1.3 |  |
| **RMM-H*^Smeg^*** | **5L38** | **1** | -3,1 | 4,9 | **-3,1** | -28,3 | 8,1 | **-27,6** | -1.1 | **-1.1** |
|  |  | **2** | -3,2 | 3,0 |  | -26,8 | 6,6 |  | -1.0 |  |

Potential local residue distortions during the MD might result in bending and tilting value over/under-estimations. Such problem might be amplified by inappropriately relaxed structural problems in starting reconfigured assemblies. Fig. S14 present an alternative view based on hexamer center of masses. For other details, please refer to Table S3.
