## Supplemental Figure S1 for "Inferring assembly-curving trends of bacterial micro-compartment shell hexamers from crystal structure arrangements"

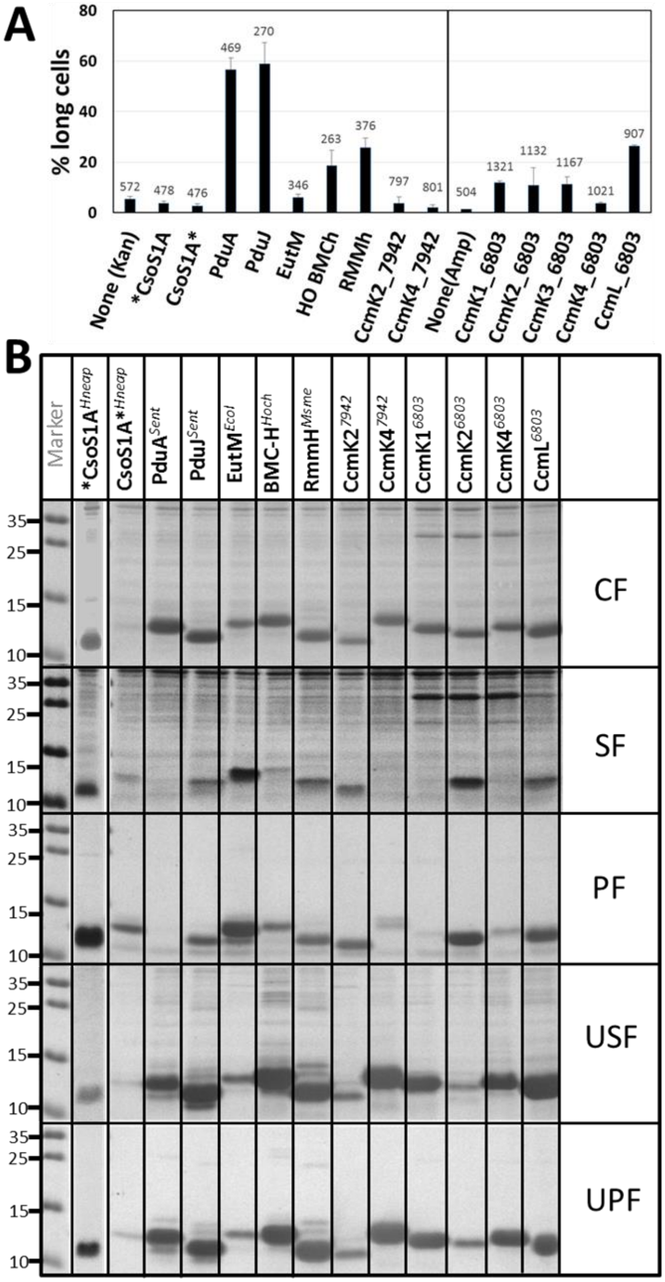


**Figure S1. Formation of potential BMC-H assemblies upon overexpression in E. coli.** *A*, Overexpression of BMC-H alters the size of cells. The percentage of BL21(DE3) longer than 1.5 times the mean size of cells carrying empty plasmids is plotted. Data were extracted from series of phase contrast microscopy images taken six hours after IPTG induction. Images were processed automatically using an adapted ImageJ macro for particle detection. The graph is divided in two portions, since expression from pET15b (*Syn6803* proteins on the right side) resulted in significantly smaller cells than from pET26b. Represented is the mean from two independent experiments (total number of cell particle counts indicated above each bar), and error bars represent standard error of the means. *B*, Coomassie-stained SDS-PAGE showing the result of the over-expression of indicated BMC-H at 37°C in autoinduced BL21(DE3) cells. From top to bottom is shown for each BMC-H: total expression level (cellular fraction, CF); soluble fractions (SF) retained in supernatants after lysis and centrifugation; purified proteins (PF) recovered from SF using standard Histag-based chromatography; urea-solubilized fraction (USF), which correspond to material remaining soluble after treatment of pelleted material in the presence of 1 M urea; urea-purified fractions (UPF), which is the material recovered from USF. Only the portion of the gel showing the proteins of interest is shown. PF and UPF fractions can be directly compared, since the two fractions were obtained after elution in the same volume and following the same procedure to load the gel. Only CcmL in UPF was 4 x lower than for all other proteins. Data for N-ter Histagged CsoS1A in the second lane (tag-position indicated by the asterisk) was from an independent experiment and is included here for comparison. As for panel *A*, *Syn6803* proteins were expressed from pET15b under ampicillin control, pET26b (kanamycin) for all others. See M&M for further details.
