## Supplemental Figure S2 for "Inferring assembly-curving trends of bacterial micro-compartment shell hexamers from crystal structure arrangements"

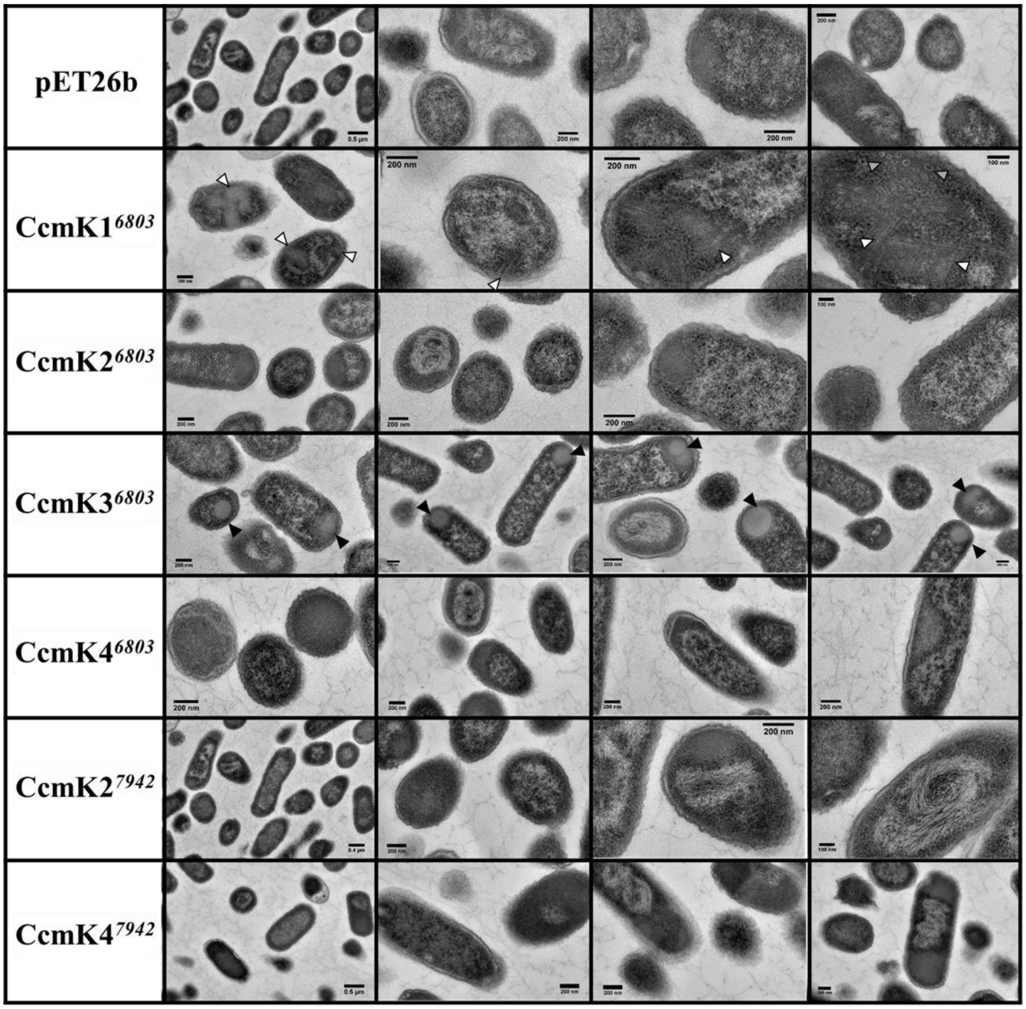


**Figure S2. Potential CcmK assembly occurrence in *E. coli* visualized by TEM.** Thin sections of bacteria over-expressing each indicated CcmK protein are presented in each lane. Only CcmK1$^{6803}$ and CcmK3$^{6803}$ produced noticeable structures, discussed in the main text. Potential 2D-layered assemblies occurring in cells expressing CcmK1$^{6803}$ are indicated by the white arrows, whereas black arrows point to CcmK3$^{6803}$ inclusion bodies. Rounded structures were also observed inside a single CcmK1$^{6803}$ cell (grey arrows). All other cases did not reveal organizations differing from what is observed in cells transformed with empty pET26b vector (first lane).
