## Supplementary figures and images for "Inferring assembly-curving trends of bacterial micro-compartment shell hexamers from crystal structure arrangements"

### Supplemental Figure S3

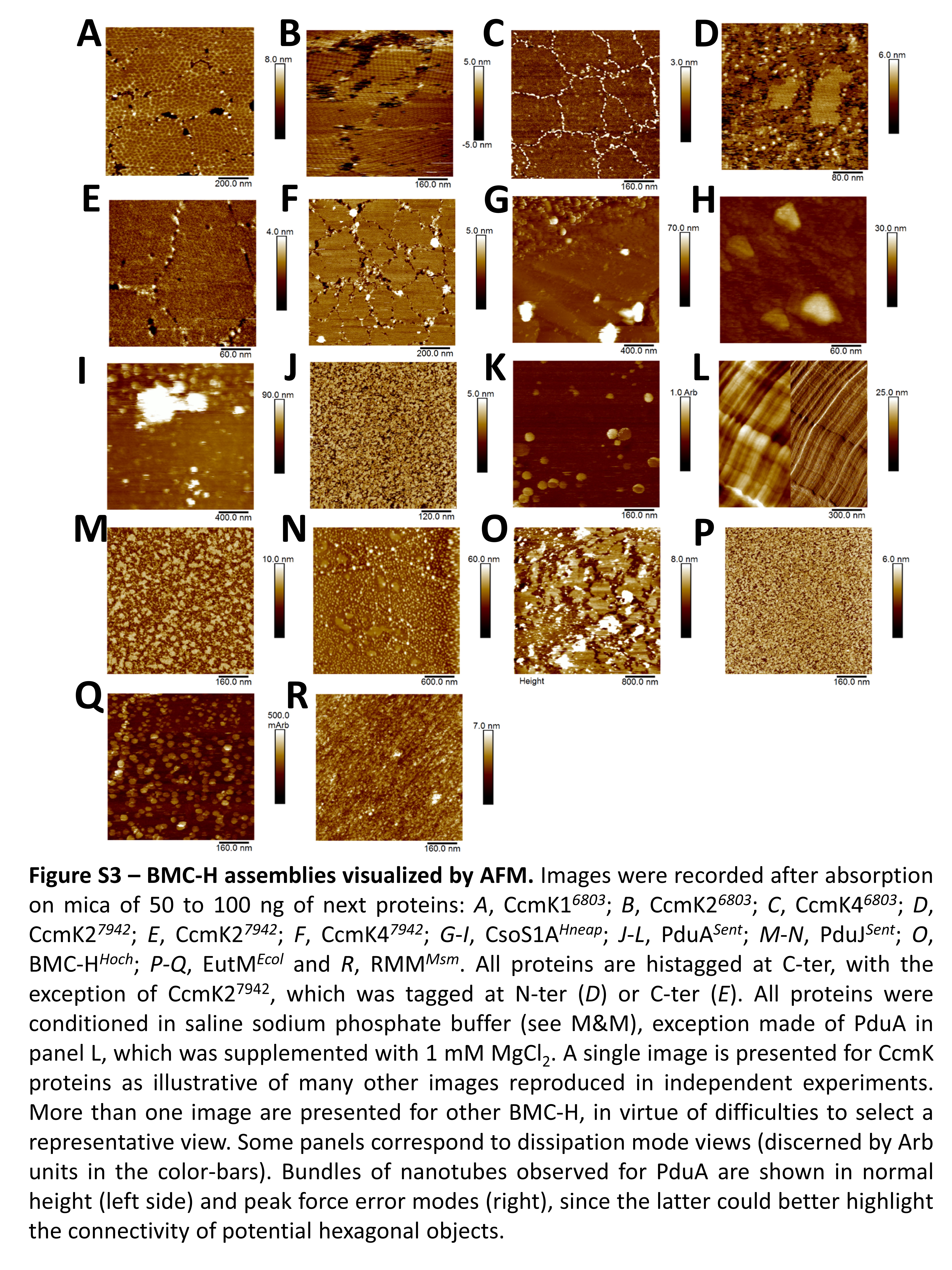

### Supplemental Figure S17

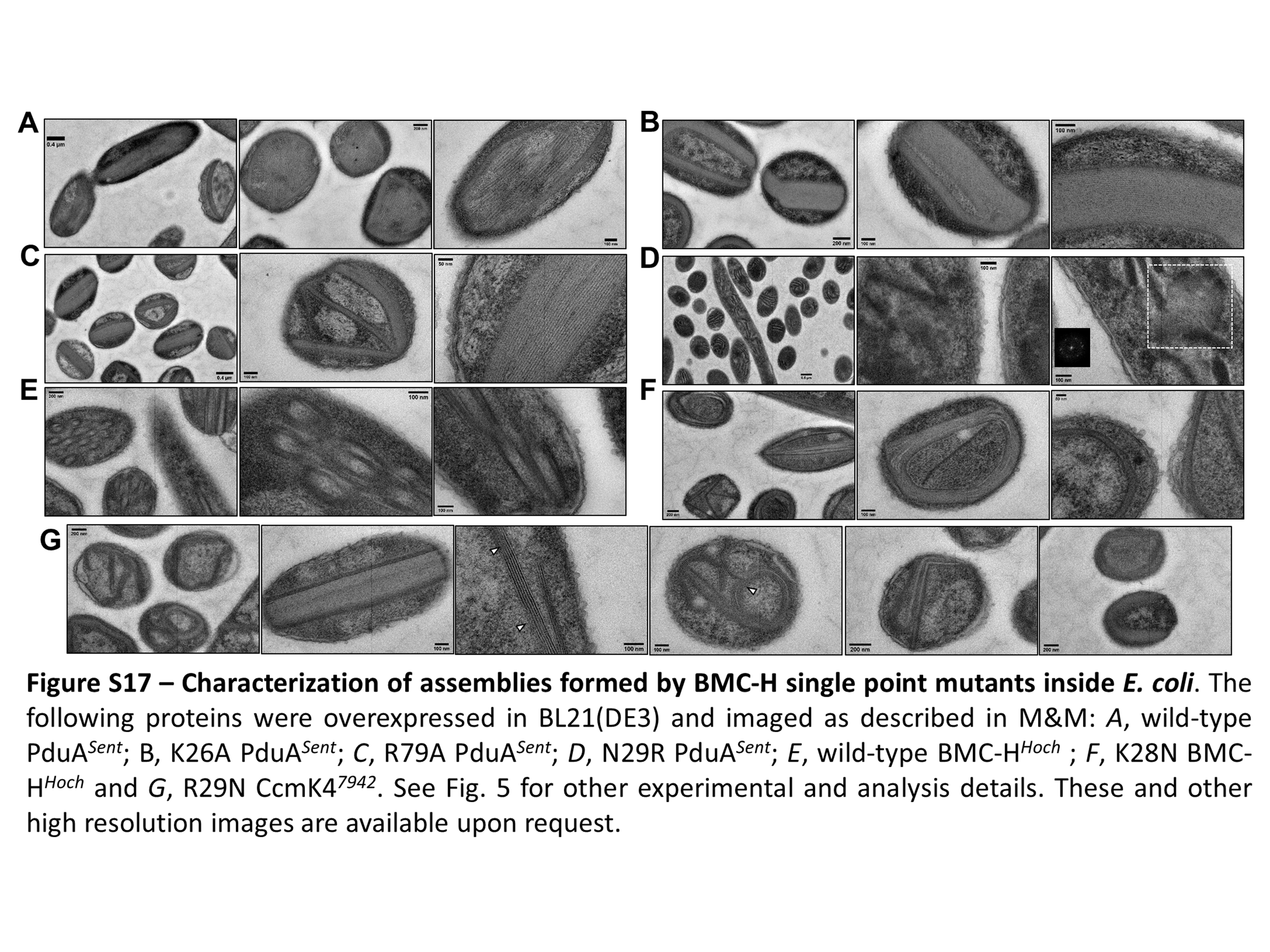
