## Supplemental Figure S4 for "Inferring assembly-curving trends of bacterial micro-compartment shell hexamers from crystal structure arrangements"

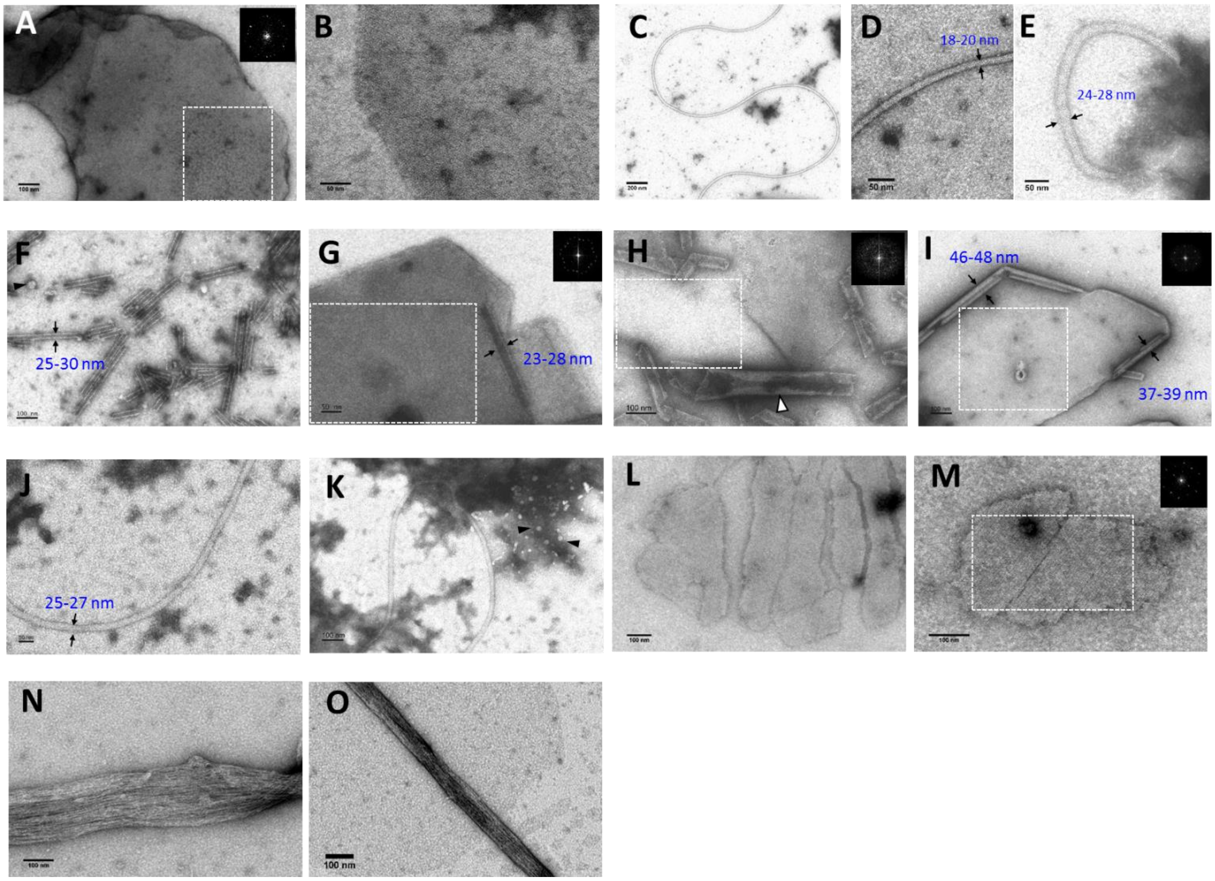


**Figure S4. BMC-H assemblies visualized by TEM.** Next proteins at 0,1-0,2 mg/mL concentrations in saline sodium phosphate buffer were imaged as described in M&M: *A-B*, CcmK1$^{6803}$; *C-E*, CsoS1A$^{Hneap}$; *F-G*, PduA$^{Sent}$; *H-I*, PduJ$^{Sent}$; *J-K*, EutM$^{Ecol}$; *L-M*, BMC-H$^{Hoch}$ and *N-O*, RMM$^{Msm}$. Structures attaching to carbon grids were recovered from drops sitting on parafilm (see M&M). Figure insets in black present the result of Fourier-transform treatments of areas outlined with white squares. Some images are illustrative of objects reproduced in independent experiments, such as nanotubes with PduA or PduJ, whereas others were less frequent (see main text). This is the case of all nano-wires and potential spheroids (indicated by black triangles), or the two fibers shown for RMM$^{Msm}$. The white triangle in panel H is to highlight a longitudinally-opened nanotube, which could be interpreted as a rolled 2D-carpet too. FFT treatments also permitted to sense 2D-organized material with next periodicities: 6,5-6,9 nm for CcmK1$^{6803}$; 6,4-6,7 nm for PduA$^{Sent}$; 6,4-6,8 nm for PduJ; 7,0-7,5 nm for BMC-H$^{Hoch}$. These and other high resolution images are available upon request for more detailed analysis.
