## Supplemental Figure S5 for "Inferring assembly-curving trends of bacterial micro-compartment shell hexamers from crystal structure arrangements"

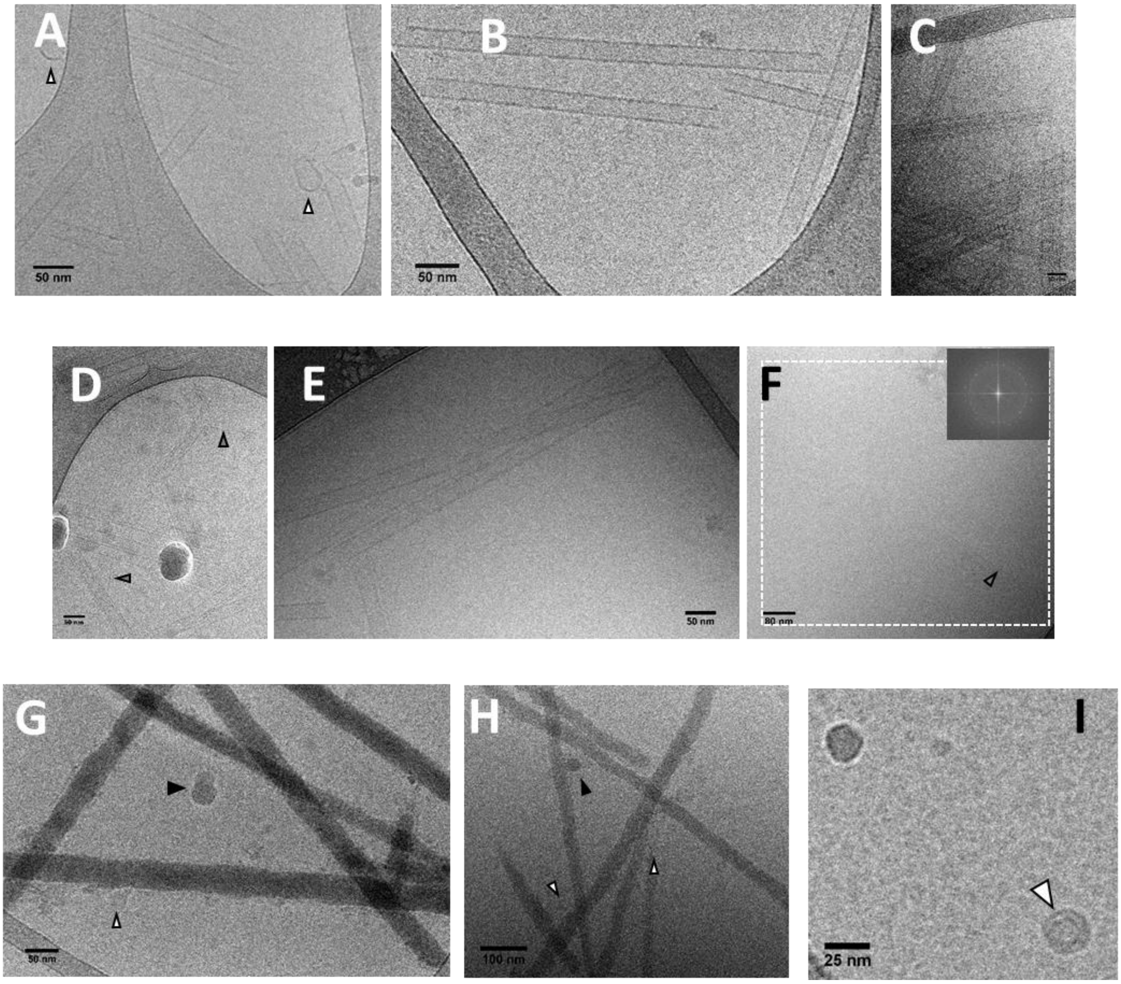


**Figure S5. Characterization of PduA, PduJ and CsoS1A assemblies by cryo-EM.** Structures were recovered on carbon grids from drops sitting on parafilm, and treated as indicated in the M&M section. Panels *A-C*, PduA$^{Sent}$; *D-F,* PduJ$^{Sent}$; *G-I*, CsoS1A$^{Hneap}$. With CsoS1A and PduA, nanotubes were the most abundant objects noticed. The relatively amorphous CsoS1A nanotubes were accompanied in occasions with spheroids of variable intensities, indicated by white and black triangles. With PduJ, faint nanotubes were detected (denser spherical particles are interpreted as cryo-EM artifacts). FT treatments of panel F or similar images also permitted to sense the presence of organized material (6,4 nm frequencies), suggestive of 2D-layered assemblies that would be too thin as to be detected visually. Edges of these 2D-assemblies might correspond to single lined contours highlighted by the gray triangles in panel D and F.
