## Supplemental Figure S6 for "Inferring assembly-curving trends of bacterial micro-compartment shell hexamers from crystal structure arrangements"

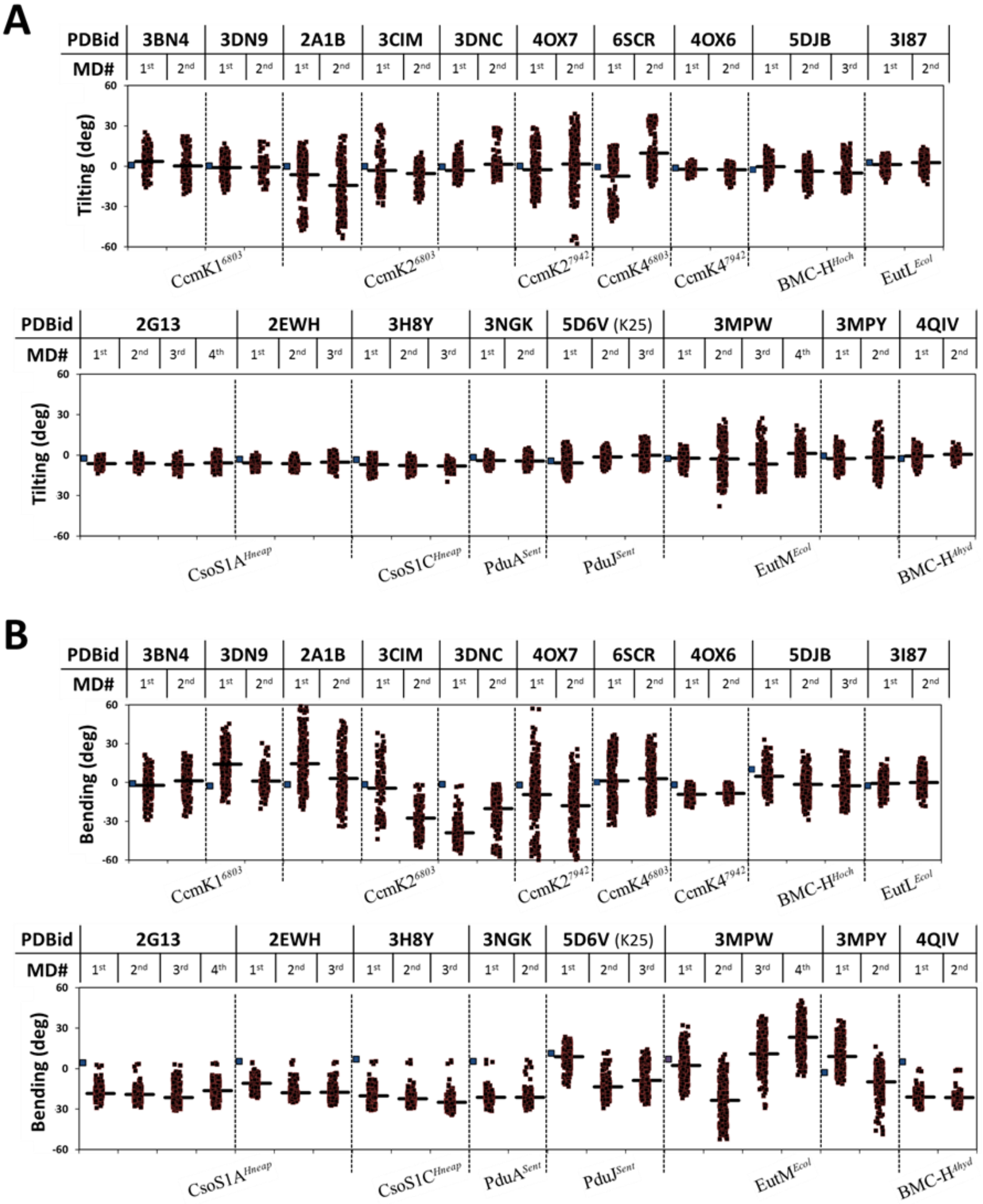


**Figure S6. BMC-H tri-hexamer behavior during MD simulations.** Plot of tilting (panel *A*) and bending (*B*) angles calculated through all-atom MD trajectories of ensembles of three BMC-H originally positioned as in crystal structures (indicated by PDBid codes on top). Tilting angles were calculated taking the Ca atom positions of M24 and Ile18 from two monomers of a given hexamer (PduA$^{Sent}$, corresponding residues in other BMC-H) with regard to same symmetric residues of the interacting hexamer counterpart. That was applied on all 3 hex-hex interfaces. Similarly bending values are based on Ca atom positions of S27 and Ile38 from one of the interfacial monomers with regard to same residues on the symmetric monomer of the opposite hexamer. Each point corresponds to one of the three measurements for a given snapshot (0.25 ns, black squares). Data from two independent 20 ns simulations are presented separately (1$^{st}$ and 2$^{nd}$). For comparison, results obtained on the EutL$^{Ecol}$ BMC-T (3I87) are presented in the last two columns of the upper portion of each panel. Thick traces represent the mean value calculated over the entire MD run. Blue squares on the most left side for each PDB case give the angle values measured for the original crystal structure. Values should be taken with caution, since occasionally readings are impacted by local distortions of the protein backbone. An alternative view is proposed by plane representations based on hexamer center of masses in Fig. 3.
