## Supplemental Figure S7 for "Inferring assembly-curving trends of bacterial micro-compartment shell hexamers from crystal structure arrangements"

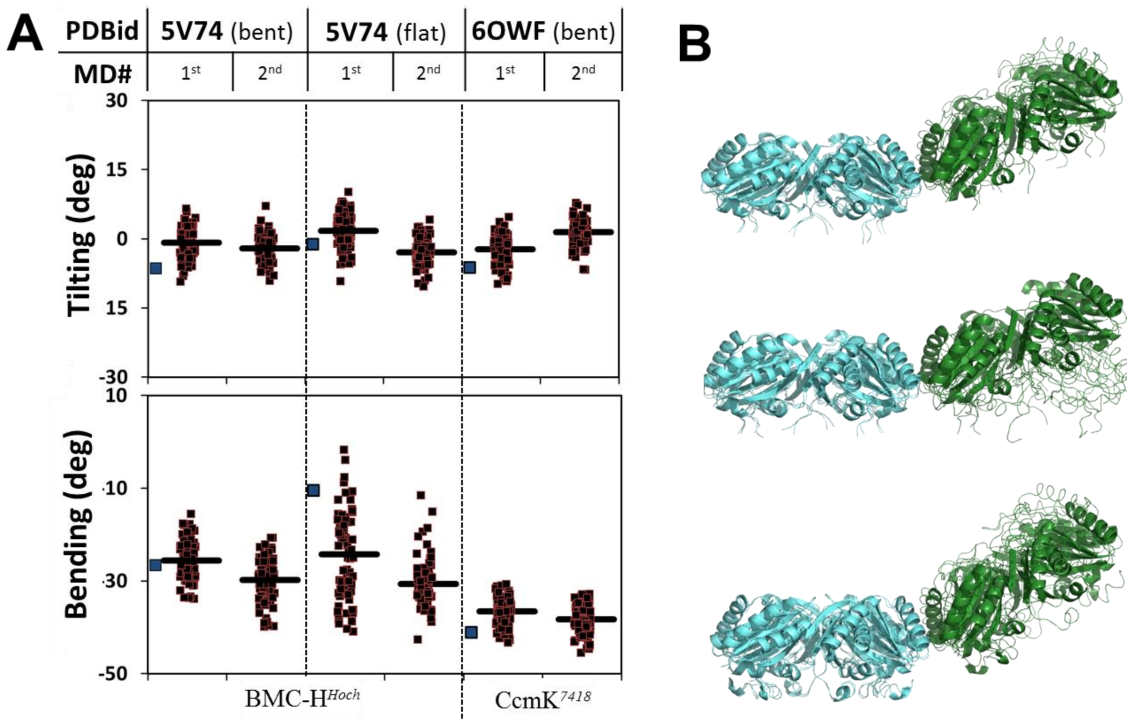


**Figure S7. MD simulations on bi-hexamers from BMC shells.** *A*, All-atom molecular dynamics of an ensemble of two hexamers extracted from published structures (PDBid codes indicated on top) and positioned in planar or curved configurations depending on the selected shell environment. See Fig. 2 for further details. *B*, Comparison of structures generated by averaging atom positions over the MD snapshots (cartoons) with the disposition in starting structures (thin traces). The two structures were superposed on backbone atom coordinates of one of the hexamers (shown in cyan, on the left side). From top to bottom: bent BMC-H$^{Hoch}$ (5V74), flat BMC-H$^{Hoch}$ (5V74) and bent CcmK$^{7418}$ (6OWF).
