## Supplemental Figure S8 for "Inferring assembly-curving trends of bacterial micro-compartment shell hexamers from crystal structure arrangements"

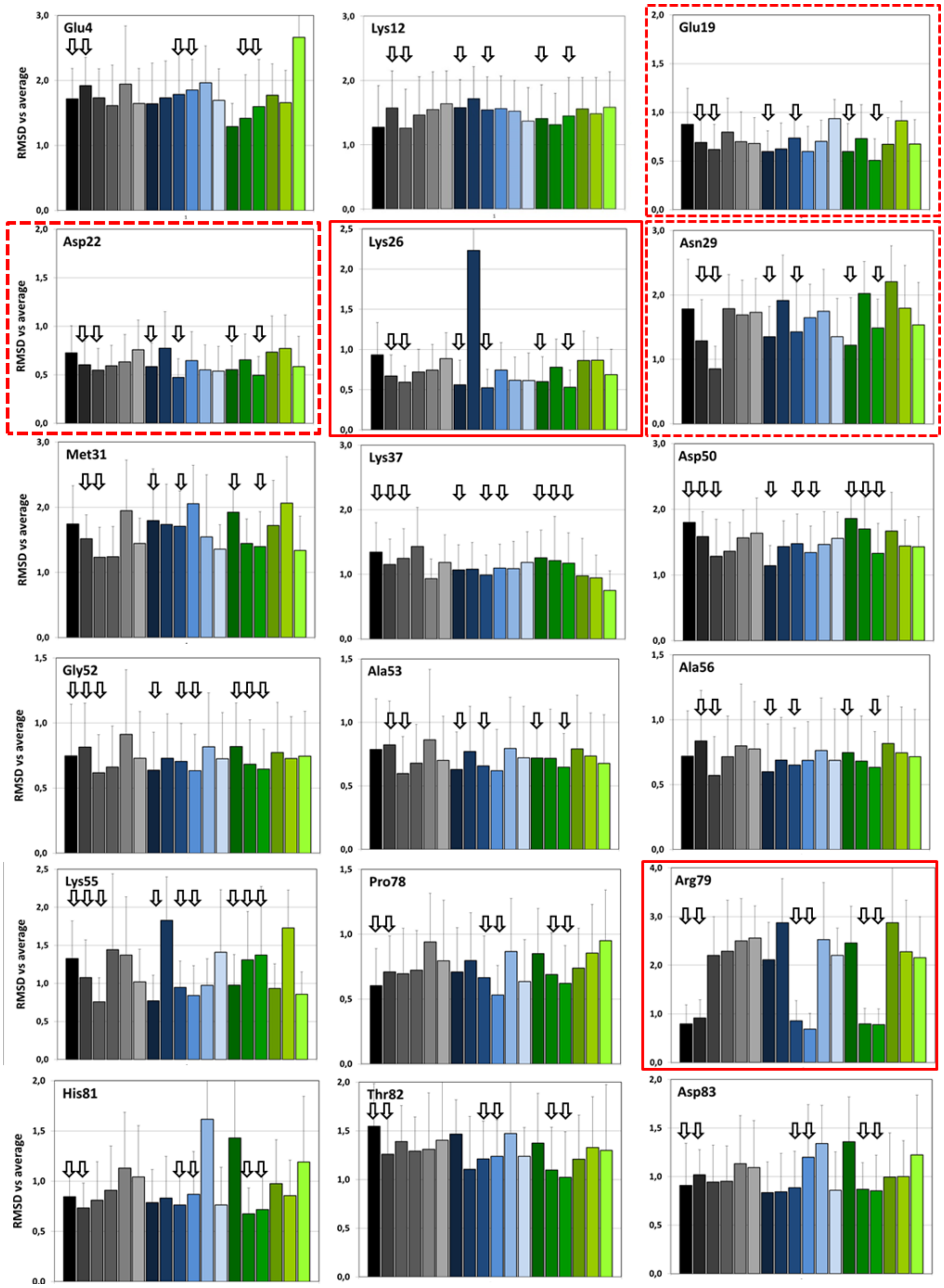


**Figure S8. RMSD evolution of PduA residues over MD simulations.** Represented is the average of root-mean-square deviations (RMSD) measured between side-chain atoms of indicated residues in each MD snapshot when compared to the residue atom coordinates in the structure averaged over the first PduA MD run. The first 8 snapshots were excluded from the calculations. Each panel present the values for a given residue in each of the 18 monomers of the tri-hexamer. Values in monomers from the first hexamer are shown in black to light grey scale, from the second hexamer with blue tonalities, green for the third. The arrows are to indicate residues from monomers that enter in contact with a neighboring hexamer. Similar results were obtained from data collected in the second MD run. Residues outlined in continuous red systematically show lower RMSD when placed at the inter-hexamer interface, in independent MD runs. Discontinuous outlines are for those residues that occur often, but not always, with lower RMSD.
