## Supplemental Figure S9 for "Inferring assembly-curving trends of bacterial micro-compartment shell hexamers from crystal structure arrangements"

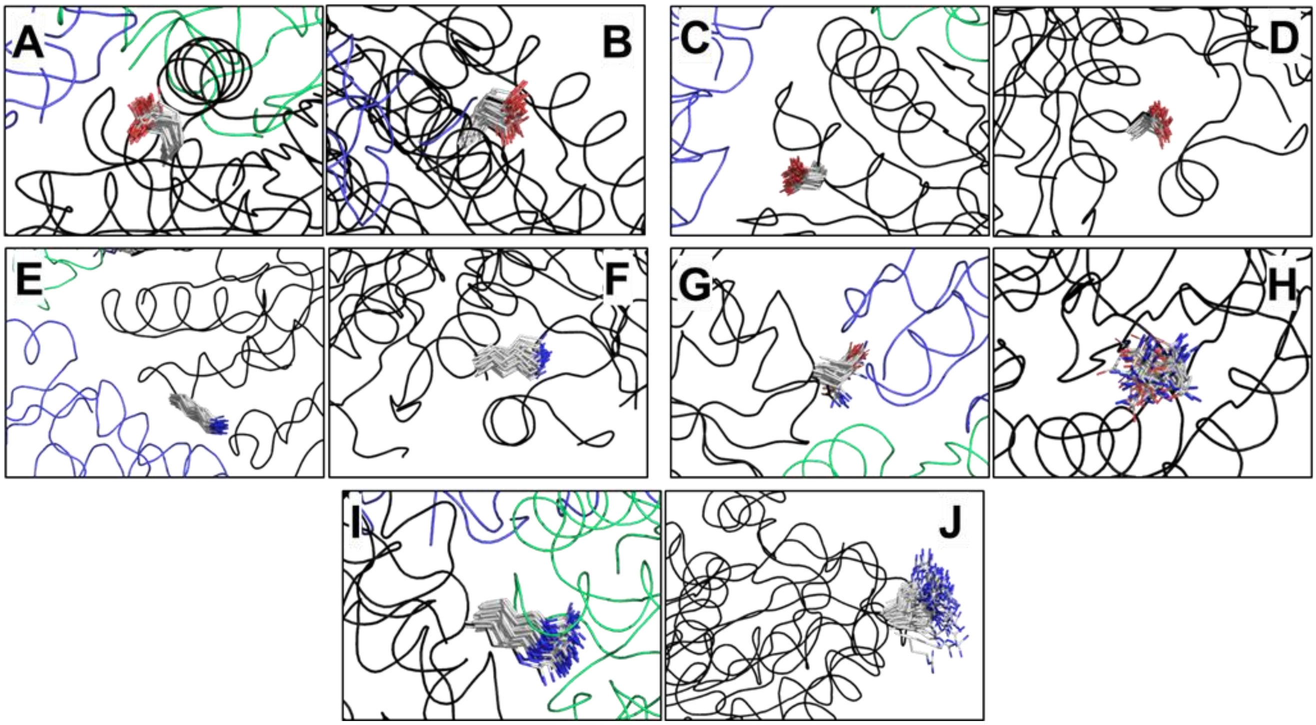


**Figure S9. Sidechain movements of selected PduA residues during MD simulations.** The view presents the side-chain conformations adopted by the several residues in the different collected snapshots of the first MD run on PduA$^{Sent}$, depending on whether the residue lies at the inter-hexamer interface (left panels) or not (right): *A-B*: Glu19; *C-D*: Asp22; *E-F*: Lys26; *G-H*: Asn29; *I-J*: Arg79. Side-chains are represented as sticks, with nitrogens blue and oxygens in red. Residues were selected from data presented in Fig. S8. All snapshot structures were superimposed on main-chain atoms of one of the hexamers (black cartoon). The two other hexamers are shown in blue or green traces.
