## Supplemental Figure S10 for "Inferring assembly-curving trends of bacterial micro-compartment shell hexamers from crystal structure arrangements"

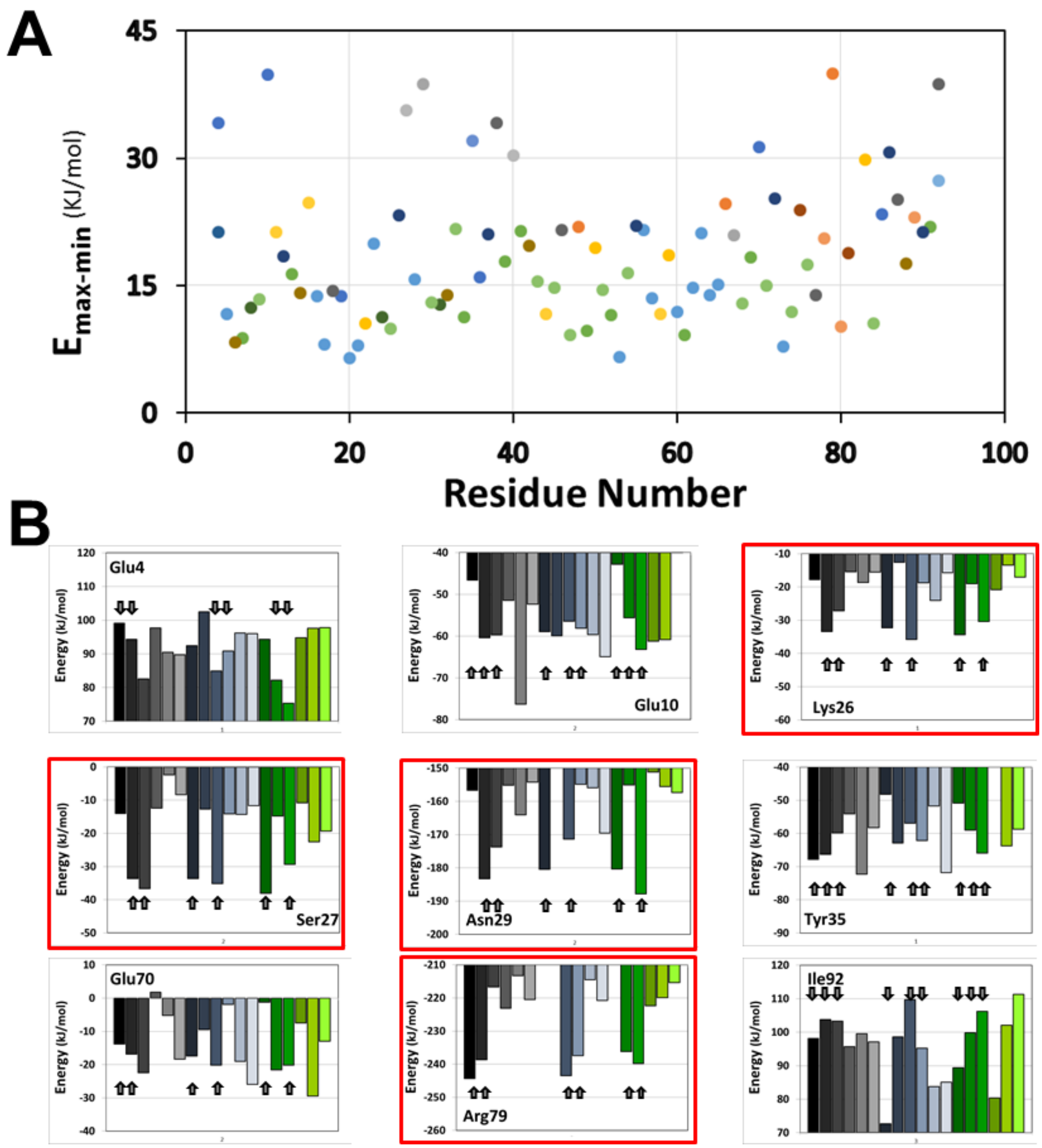


**Figure S10. Energetic contribution of PduA residues to the stabilization of the bent assembly.** *A*, Interval of energies contributed by every residue of PduA, when comparing the 18 monomers of the tri-hexamer assembly. The ordinate presents the energy interval measured between the less and most stabilizing position. The most similar (lowest RMSD) snapshot to the averaged structure of the first MD run was selected for the analysis. Energy computation was done with GROMOS96 implemented in Swiss-PDBViewer. *B*, Estimated energy contribution of selected residues in the 18 different emplacements of the trihexamer. Only a few residues among those analyzed are presented. Data are colored as in Fig. S8. The arrows are to identify residues in monomers that interact with the neighbor hexamer. Similar results were obtained from data collected in the second MD run. Outlined in red are residues that systematically contribute lowest energies when positioned at the inter-hexamer interface, for both MD runs.
