## Supplemental Figure S11 for "Inferring assembly-curving trends of bacterial micro-compartment shell hexamers from crystal structure arrangements"

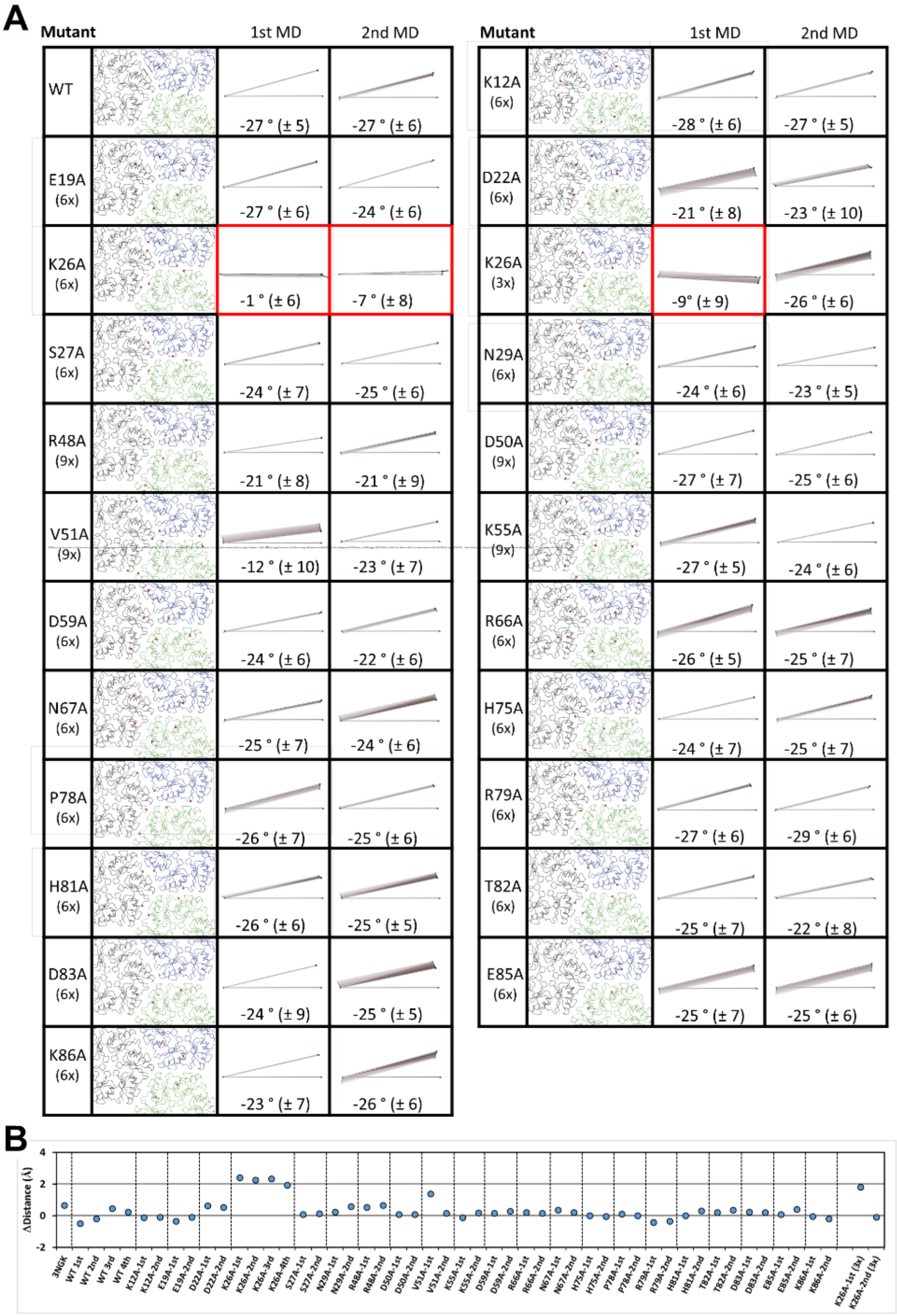


**Figure S11. MD behavior of PduA tri-hexamer assemblies with selected residues mutated into alanine.** *A*, Residues indicated in the first lane were replaced by alanine in the 6 or 9 monomers of the trihexamer assembly that lie at the interface. An assembly with only three K26 positions mutated was also simulated. The Cα of such residues is indicated by red spheres in the second column. The result of two MD runs is presented following plane representations like those of Fig. 3. Indicated bending angle values were evaluated like in Table S3. Please notice that local distortions caused during the MD might in occasions result in under/overestimations. In the case of the K26A mutant (6x, outlined in red), four MD runs were carried out, with similar qualitative results. *B*, Effect of mutation on the distance between hexamers during the MD run. In the ordinate axes is represented the difference between the averaged distance calculated for the three hexamers (center of masses) in the averaged structure of a given MD simulation, and the average distance calculated from four independent MD runs on wild-type (WT) PduA, which are shown on the most left side, after the value measured for the PduA crytal (3NGK). Data obtained in independent MD run repetitions are denoted by 1st and 2nd label extensions below the X-axis.
