## Supplemental Figure S12 for "Inferring assembly-curving trends of bacterial micro-compartment shell hexamers from crystal structure arrangements"

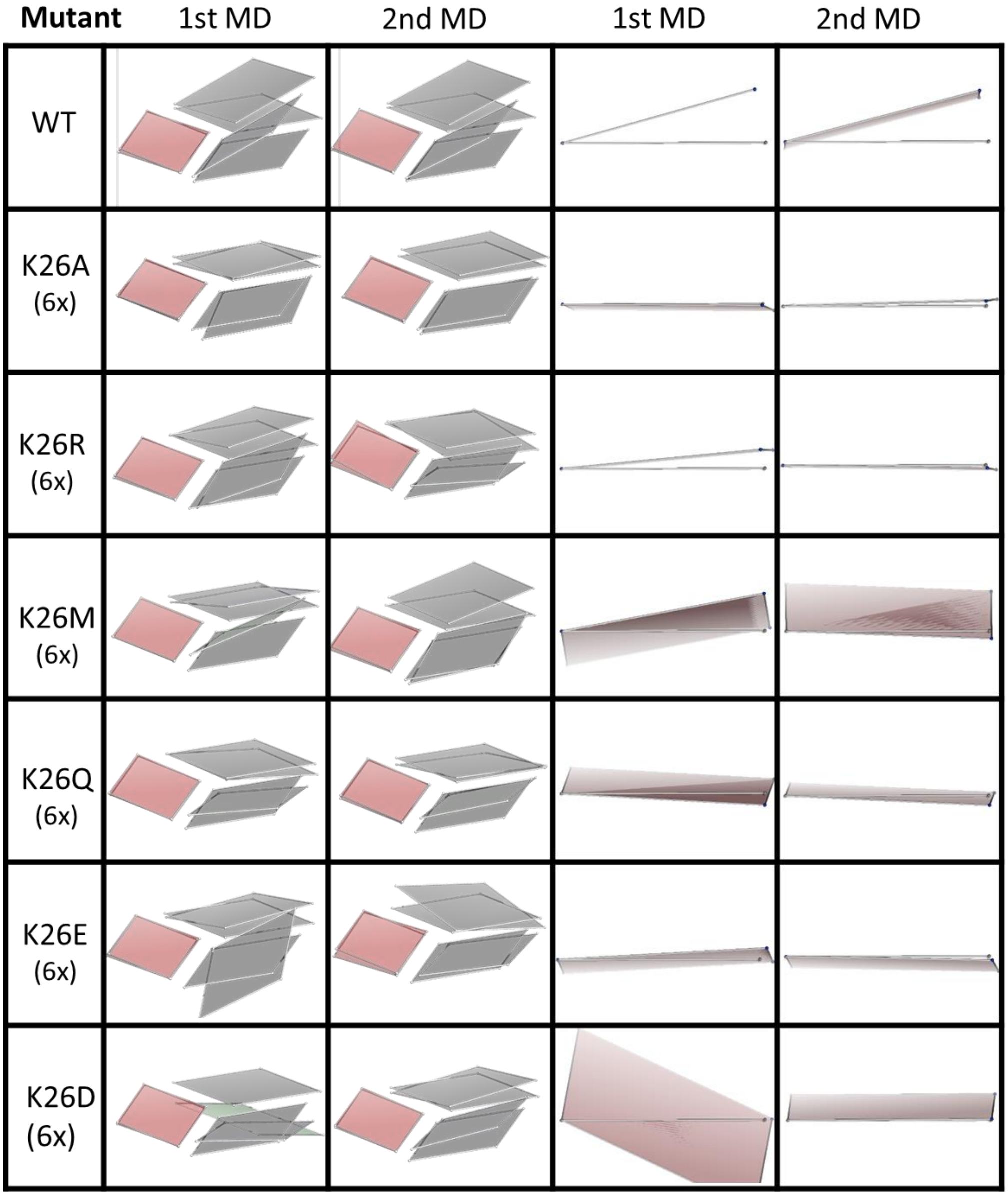


**Figure S12. MD consequences of replacement of K26 of PduA by other residue types**. Lys26 was replaced by residues indicated in the first column in the 6 monomers located at the contacting interface between subunits. The result of two MD runs is presented following plane representations based on center of mass of hexamers main-chain atoms like in Fig. 3 (right side). An alternative view is given on the left side, with each hexamer of the starting tri-hexamer assembly or of the MD average prepared taking the Ca positions of V53 (CcmK1*6803*, corresponding residues in other BMC-H) from the three monomers of each hexamer that are in contact with other hexamers.
