## Supplemental Figure S13 for "Inferring assembly-curving trends of bacterial micro-compartment shell hexamers from crystal structure arrangements"

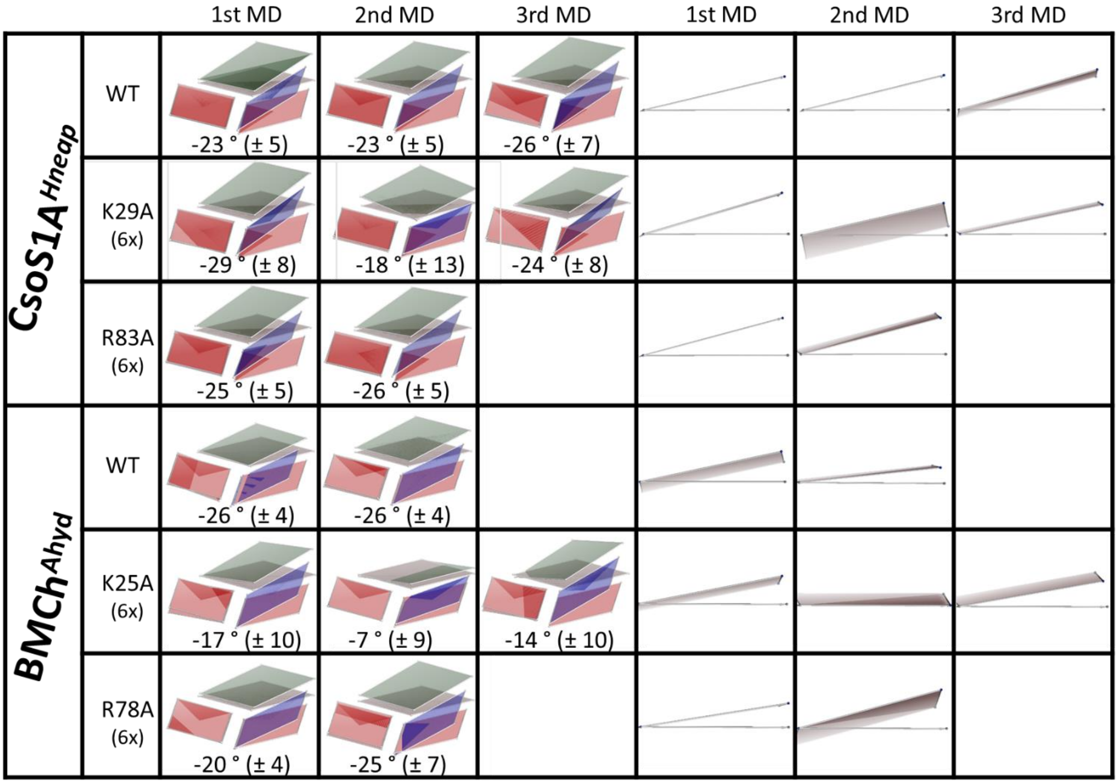


**Figure S13. MD behavior of *Ass-A* tri-hexamer BMC-H assemblies with interfacial Lys and Arg residues mutated into alanine.** Key interfacial Lys and Arg, residues indicated in the second column, were replaced in CsoS1A$^{Hneap}$ (2G13) or BMC-H$^{Ahyd}$ (4QIV) by alanine in the 6 monomers of the tri-hexamer assembly that lie at the interface. Data from several independent MD runs are presented. Data for wild-type versions are shown for the ease of comparison. Hexamers are either represented by planes prepared from positions of main-chain atoms like those presented in Fig. S12, or from the center of mass of each hexamer, likewise in Fig. 3. Please notice that indicated average bending angles, calculated as for Table S3, can be under/overestimations in virtue of local alterations caused by the presence of mutated positions.
