## Supplemental Figure S14 for "Inferring assembly-curving trends of bacterial micro-compartment shell hexamers from crystal structure arrangements"

| BMC-H | PDB id | 1st MD | 2nd MD | BMC-H | PDB id | 1st MD | 2nd MD |
|---|---|---|---|---|---|---|---|
| CcmK1$^{6803}$ | 3BN4 | 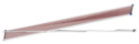 | 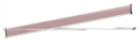 | CcmK4$^{7942}$ | 4OX6 | 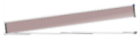 | 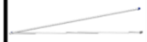 |
| CcmK2$^{6803}$ | 2A1B | 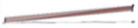 | 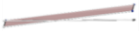 | EutM$^{Ecol}$ | 3MPW | 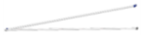 | 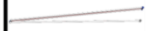 |
| CcmK2$^{7942}$ | 4OX7 | 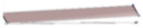 | 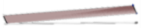 | BMCh$^{Hoch}$ | 5DLB | 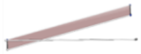 | 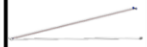 |
| CcmK4$^{6803}$ | 6SCR |  | 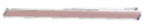 | RMM-H$^{Msme}$ | 5L38 | 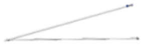 | 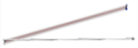 |
| CcmK4$^{6803}$ *pentamutant* | 6SCR | 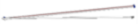 | 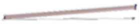 | | | | |

**Figure S14. Dynamic behavior of tri-hexamers reconfigured in *Ass-A* disposition.** Comparison of the average structure generated for all snapshots of each MD simulation with the disposition at time 0. Other details are like for Fig. 3. The CcmK4$^{6803}$ penta-mutant carried the next changes with regard to the WT version: R30N (6x), Q53G (9x), E54A (9x), E85T (6x) and N86D (6x).
