## Supplemental Figure S15 for "Inferring assembly-curving trends of bacterial micro-compartment shell hexamers from crystal structure arrangements"

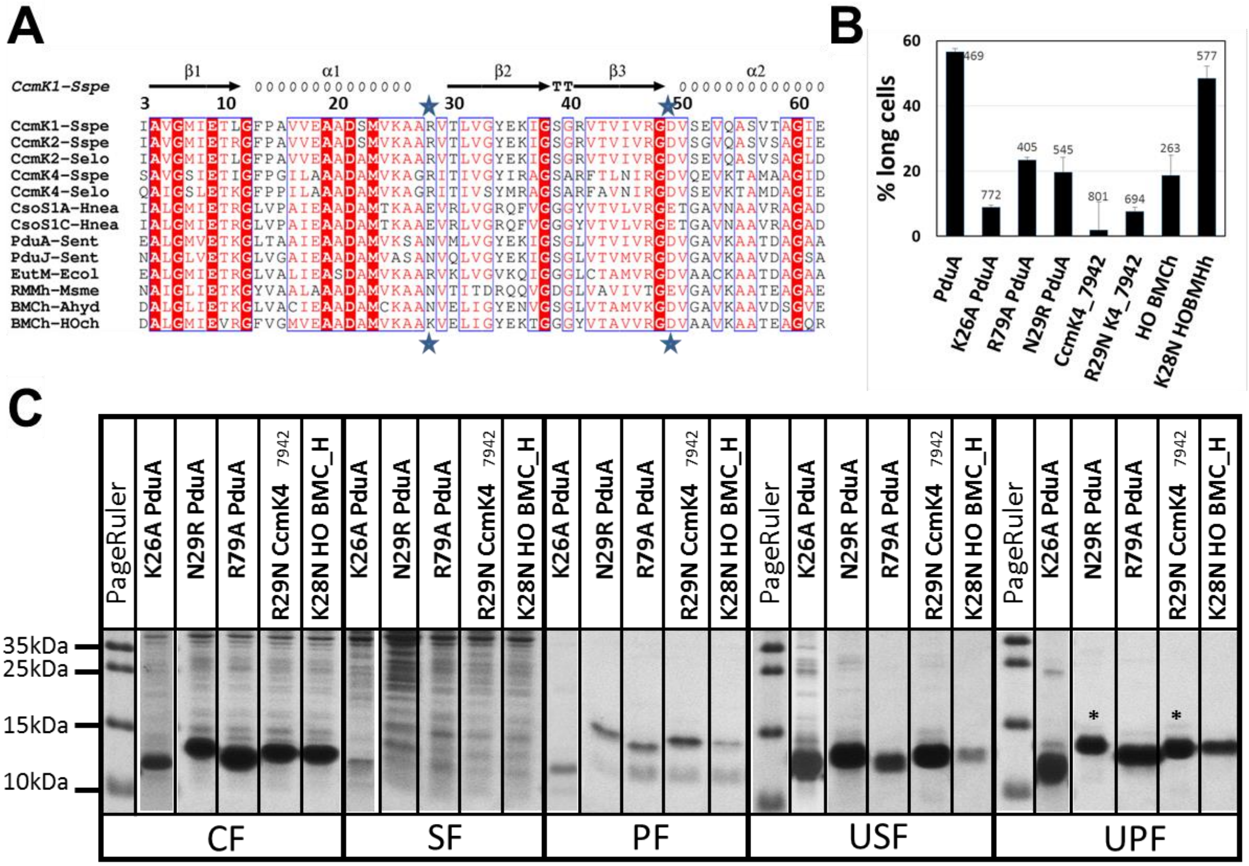


**Figure S15. Assembly of BMC-H with key interfacial residue mutations.** *A*, sequence alignments indicate the existence of potential ionic-pairing residues in a subset of BMC-H. Data from herein studied PDB hits were recovered from the RCSB databank. Extensions at N-terminus, including purification tags, and last 30 to 40 residues were removed to facilitate visual comparisons. Alignment was carried out with Clustal Omega software, and the final figure prepared online with ESPript. Secondary structural elements and residue numbering shown on top are from CcmK1$^{6803}$ 3BN4 structure (Sspe and Selo denote *Syn. sp. PCC 6803* and *Syn. elongatus PCC 7942*). Stars indicate positions implicated in ionic interactions in non-*Ass-A* is assemblies. *B-C*, Effect of single point mutations on the percentage of elongated cells (*B*) and *o*n recovered protein yields (*C*) when BL21(DE3) strains were induced with IPTG and treated exactly as for experiments of Fig. S1. Amounts in the 3$^{rd}$ and 5$^{th}$ lanes of the UPF gel (indicated by asterisk) should be multiplied by 4, since sample load was 4 times lower than with other samples.
