## Supplemental Figure S16 for "Inferring assembly-curving trends of bacterial micro-compartment shell hexamers from crystal structure arrangements"

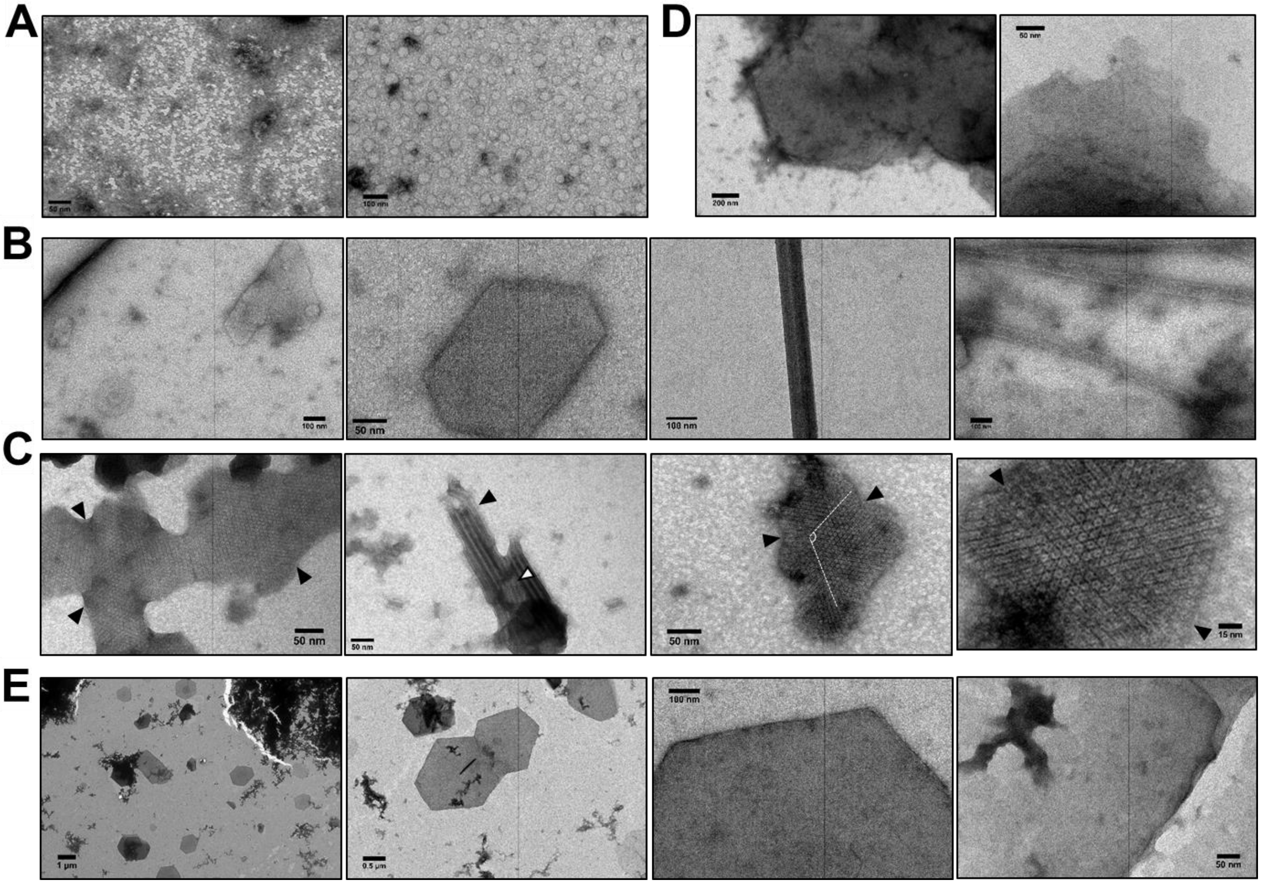


**Figure S16. Alteration of BMC-H assembly by single point mutations of interfacial residues.** TEM images collected at 0,1-0,2 mg/mL concentrations of purified proteins in saline sodium phosphate buffer: *A*, K26A PduA$^{Sent}$; *B*, R79A PduA$^{Sent}$; *C*, N29R PduA$^{Sent}$; *D*, K28N BMC-H$^{Hoch}$ and *E*, R29N CcmK4$^{7942}$. *A*, only small patches were detected with the K26A PduA mutant. Imaging was hampered by the presence of abundant skinny material. The patches could not be unambiguously assigned to protein assemblies. *B*, a few 2D patches and a single nanotube were observed in experiments with R79A PduA. In occasions, nanotube like structures were imaged, but might correspond to skin wrinkling. *C*, 2D-assemblies with well-defined hexamers were abundant with the N29R mutant. Inter-hexamer separations (7 to 9 Å) were compatible with *Ass-B* organizations. Striped arrangements were disposed with 117 to 125° angles, as indicated in the third image. Potential nanotubes were also observed for this mutant, always appearing as bundles. Mixed patches like those pointed by the arrows in the second image could reflect fluctuations between flat to bent organizations. The K28N BMC-H$^{Hoch}$ (*D*) and R29N CcmK4$^{7942}$ (*E*) mutants gave rise to compact 2D-assemblies, without evidence of formation of curved structures. These and other high resolution images are available upon request for more detailed analysis.
